# The human-specific lncRNA HD-LINC drives lipid droplet-associated neurodegeneration in Huntington’s disease

**DOI:** 10.64898/2026.10.23.761349

**Authors:** Ji-Sun Kwon, Liyuan Wang, Naga R. Ganti, Juncheng Li, Shengnan Xia, Osama Al-Dalahmah, Andrew S. Yoo

## Abstract

Huntington’s disease (HD) is characterized by degeneration of striatal medium spiny neurons (MSNs), yet the human-specific regulatory mechanisms that contribute to neuronal vulnerability remain poorly understood. Here, we identify HD-LINC (LINC01139), a human-specific long intergenic noncoding RNA selectively induced in symptomatic HD patient-derived MSNs (HD-MSNs) and human HD caudate. Single-cell transcriptomic analysis in HD-MSNs revealed that HD-LINC was enriched in a symptomatic HD neuronal state associated with lipid metabolic and protein clearance regulation, while not observed in non-degenerative reprogrammed MSNs derived from pre-manifest HD individuals. Reducing HD-LINC attenuated mutant HTT aggregation, dystrophic neurites, DNA damage, and neuronal death, whereas HD-LINC overexpression exacerbated these phenotypes. Transcriptomic and lipidomic analyses further implicated HD-LINC in neuronal lipid regulation. We found that HD-MSNs exhibited aberrant accumulation of enlarged, low-mobility lipid droplets (LDs) within neurites, a previously unrecognized phenotype in patient-derived neurons. Critically, increased lipid peroxidation was observed within LD-enriched dystrophic neurites in HD-MSNs. HD-LINC knockdown cleared neuritic LD accumulation, restored soma-enriched LD distribution, and reduced lipid peroxidation. Finally, we developed antisense oligonucleotides that effectively reduced HD-LINC and reversed HD-LINC-associated phenotypes, nominating HD-LINC lowering as a potential therapeutic strategy in HD. These results identify HD-LINC as a human-specific regulator of neuronal LD spatial organization, lipid peroxidation, and neuronal degeneration in HD. They further reveal dysregulated neuritic LD accumulation driven by HD-LINC, and highlight lincRNA-mediated lipid dysregulation as a significant contributor to neuronal vulnerability to degeneration.

## Introduction

Huntington’s disease (HD) is a dominantly inherited neurodegenerative disorder caused by a CAG repeat expansion in the HTT gene^1^. Because mutant HTT is ubiquitously expressed across diverse cell types, the molecular basis underlying the selective vulnerability of striatal medium spiny neurons (MSNs) remains a central unresolved question in HD^2^. Transcriptomic analyses of HD patient brains and animal models have revealed widespread gene-expression alterations, largely focused on protein-coding genes^3–5^. However, the contribution of noncoding regulatory mechanisms to the pronounced cell-type-specific degeneration observed in human MSNs remains incompletely understood. Human-specific and cell-type-selective noncoding RNAs represent a compelling regulatory layer that could provide insights into mechanisms governing neuronal vulnerability in HD^6,7^.

Studying such human-specific noncoding mechanisms underlying adult-onset neurodegeneration requires an experimental system that preserves age-associated transcriptional and epigenetic programs. Direct neuronal reprogramming using microRNAs, miR-9/9* and miR-124 (miR-9/9*-124), in combination with MSN lineage factors, generates neurons that maintain donor epigenetic age, transcriptional aging signatures, and age-associated microRNA landscapes^8,9^. Unlike iPSC-derived neurons, whose age-associated molecular signatures are reset during pluripotency induction^10^, directly induced MSNs from HD patients retain age-associated regulatory features and recapitulate HD-associated neurodegenerative phenotypes, including mutant HTT (mHTT) aggregation, DNA damage, and neuronal death^11,12^. These features make them well suited for identifying non-conserved drivers of vulnerability, including noncoding RNAs.

Growing evidence implicates lipid dysregulation in HD. Alterations in cholesterol metabolism and fatty-acid homeostasis have been reported in HD patient brains and model systems^13–15^, and increased lipid-droplet (LD) accumulation has also been observed^16,17^. However, how LD accumulation is regulated, how LDs are organized within neurons, and how these features relate to neuronal dysfunction remain poorly understood. Beyond their established roles in transcriptional regulation, long noncoding RNAs (lncRNAs) have emerged as critical regulators of lipid metabolism and LD biology in multiple non-neuronal systems^18,19^. Whether lncRNAs, specifically human-specific lncRNAs, regulate neuronal lipid homeostasis and the spatial organization of LDs, and how these processes contribute to the degeneration of human MSNs in HD, remain largely unexplored.

Here, we identify LINC01139, a human-specific lincRNA (96.6%, 38.9%, and 0% sequence identity with chimpanzee LOC744344 exons 1–3, respectively), as a regulator of neuronal vulnerability in HD-MSNs. We term this transcript HD-induced lincRNA (HD-LINC), as it is readily detected in HD-MSNs derived from symptomatic HD individuals as well as in human HD caudate tissue, while not detected in neurons generated from premanifest individuals. While LINC01139, previously termed LINK-A in the cancer context, has been characterized for its regulation of HIF1α and AKT signaling^20,21^, its role in HD neurodegeneration remains unknown. Single-cell transcriptomic analysis revealed that HD-LINC is enriched in a symptomatic HD neuronal state associated with lipid metabolism and protein clearance regulation. Functionally, HD-LINC promotes dysregulated neuritic LD accumulation, whereas its repression reverses this neuritic LD accumulation and restores a control-like soma-enriched LD distribution. HD-LINC also promotes lipid peroxidation associated with neuritic degeneration and mHTT aggregation. HD-LINC repression further attenuates dystrophic neurites and neuronal death by lipid-peroxidation. Antisense oligonucleotide (ASO)-mediated repression of HD-LINC recapitulates these protective effects. Together, these findings identify HD-LINC as a human-specific pathogenic driver of LD accumulation in dystrophic neurites, contributing to neuronal vulnerability in HD.

## Results

### Symptomatic HD-MSNs exhibit a distinct neuronal state associated with lipid metabolism and protein clearance

To generate age-maintained directly reprogrammed MSNs^9^, we used the established microRNA (miR-9/9*-124)-based direct neuronal reprogramming protocol, in which miR-9/9*-124 drives the neurogenic conversion of adult human fibroblasts, while transcription factors, CTIP2, DLX1/2, and MYT1L guide the conversion toward the MSN lineage^8^ (Fig. 1A). Because MSNs directly reprogrammed from fibroblasts obtained from symptomatic HD patients (HD-MSNs) show greater neuronal death compared to reprogrammed MSNs from young, presymptomatic HTT mutation carriers (pre-HD-MSNs)^11,12^, we first investigated transcriptional changes in single cells associated with disease stage using single-cell RNA-seq. While both groups contained four transcriptional clusters, the proportion of cells in cluster 2 was substantially higher in symptomatic HD-MSNs than pre-HD-MSNs (Fig. 1B and Supplementary Fig. 1A). Because both pre-HD-MSNs and HD-MSNs carry mutant HTT, this disease-stage-associated shift suggested that cluster 2 represents a transcriptional state associated with symptomatic progression rather than simply the presence of the HD mutation. To further characterize this symptomatic HD-associated state, we examined genes with established relevance to HD pathology and neuronal stress. TSPO, an outer mitochondrial membrane protein upregulated in a range of cellular stress and neuroinflammatory conditions^22^, showed increased expression in cluster 2. HMOX1, an oxidative stress-responsive heme oxygenase associated with ferroptotic stress^23,24^, and DDIT4, a stress-inducible inhibitor of mTORC1 signaling implicated in mHTT-induced neuronal toxicity^25,26^, were also elevated, together with CASP6, a protease implicated in pathogenic HTT cleavage and HD-associated neuronal degeneration^27^ (Fig. 1C). These findings indicate that cluster 2 is marked by genes associated with neuroinflammation, oxidative stress, stress response, and proteolytic processes.

**Figure 1.**
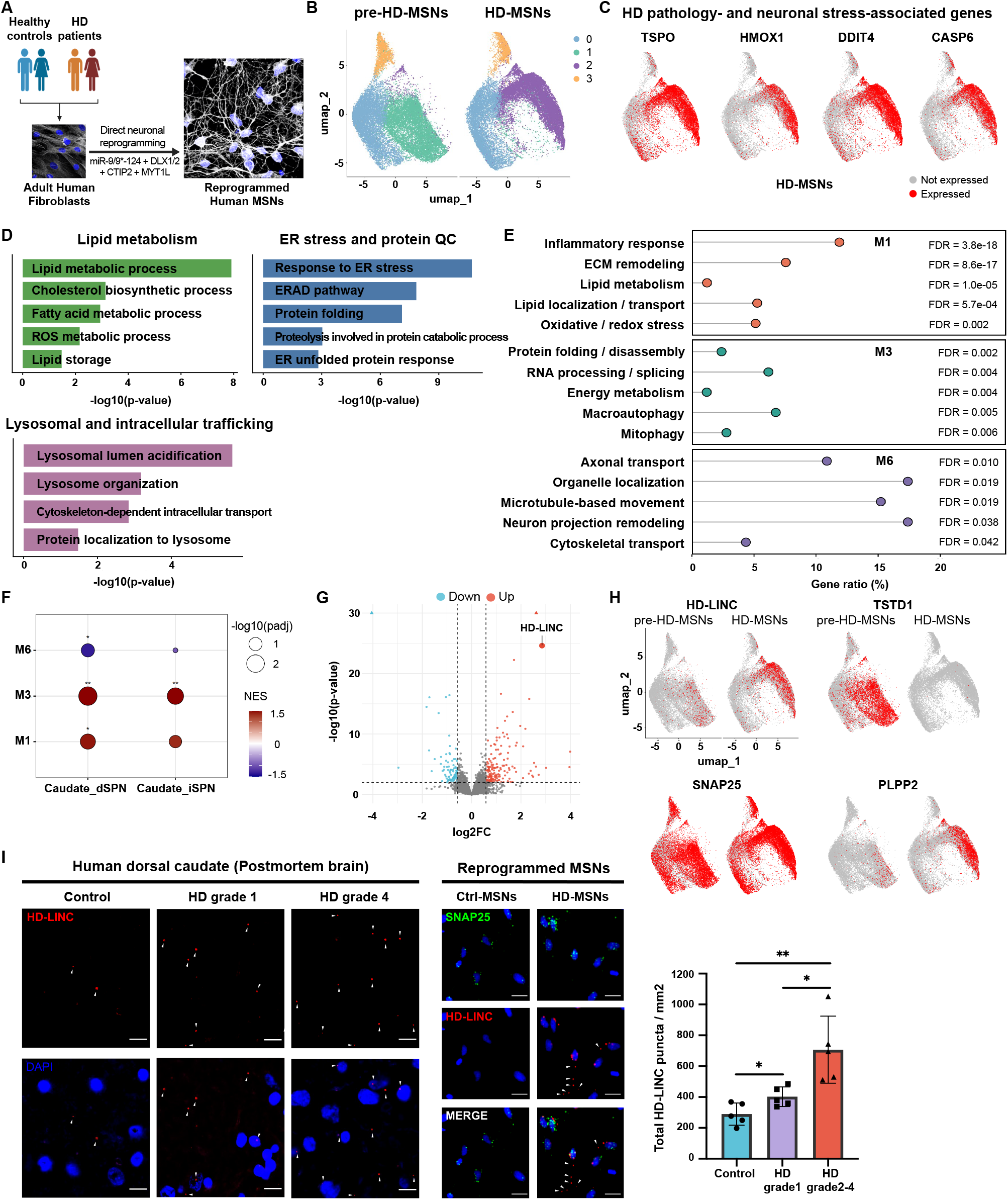
HD-LINC is selectively induced in symptomatic HD-MSNs and human HD caudate. **(A)** Experimental workflow for generating human medium spiny neurons (MSNs) by direct reprogramming of adult fibroblasts derived from healthy control individuals and HD patients using miR-9/9*–124 together with DLX1/2, CTIP2, and MYT1L. **(B)** UMAP visualizations showing transcriptional clusters in pre-symptomatic and post-symptomatic HD-MSNs. Clusters are indicated by color. **(C)** Feature plots showing expression of representative HD pathology- and neuronal stress-associated genes, including TSPO, HMOX1, DDIT4, and CASP6. Red, expressed; grey, not expressed. **(D)** Gene ontology (GO) terms associated with genes upregulated in the HD-LINC-enriched cluster 2 relative to other transcriptional clusters in symptomatic HD-MSNs, highlighting lipid metabolism, ER stress and proteolysis, and lysosomal and intracellular trafficking pathways. Bars represent −log_10_ (P value). **(E)** Functional enrichment analysis of hdWGCNA modules M3, M1, and M6 identified by comparing the HD-LINC-enriched cluster 2 with other transcriptional clusters in symptomatic HD-MSNs. Selected enriched biological processes, gene ratios, and false discovery rates (FDRs) are shown. **(F)** Pre-ranked GSEA of HD-LINC-associated hdWGCNA modules in postmortem human caudate dSPNs and iSPNs using HD-versus-control differential expression rankings (Paryani et al.). Dot color indicates NES; dot size indicates −log_10_ (adjusted P value). Benjamini–Hochberg adjustment was applied within each cell type. *Adjusted P < 0.05, **P < 0.01, ***P < 0.001. **(G)** Volcano plot of differentially expressed protein-coding genes and lncRNAs between directly reprogrammed Ctrl-MSNs and HD-MSNs. HD-LINC (LINC01139) is highlighted. Blue, downregulated genes; red, upregulated genes. **(H)** Feature plots showing expression of HD-LINC, TSTD1, SNAP25, and PLPP2 in pre-symptomatic and post-symptomatic HD-MSNs. Red, expressed; grey, not expressed. **(I)** Left, representative RNAscope images of HD-LINC signals in postmortem human dorsal caudate from control, HD grade 1, and HD grade 4 samples. Middle, representative RNAscope images of directly reprogrammed Ctrl-MSNs and HD-MSNs showing HD-LINC signals (red), SNAP25 (green), and DAPI (blue). Right, quantification of total HD-LINC puncta per mm² in MAP2-positive neurons from control, HD grade 1, and HD grade 2–4 caudate samples. For the quantification in I, each dot represents one independent postmortem human caudate sample; n = 5 samples per group. Data are mean ± s.e.m. Statistical significance in I was determined by one-way ANOVA followed by Tukey’s multiple-comparisons test. *P < 0.05, **P < 0.01. Scale bars, 10 μm (postmortem caudate) and 20 μm (repro-grammed MSNs).

Gene ontology (GO) analysis of genes enriched in cluster 2 relative to clusters 0, 1, and 3 further identified pathways related to lipid metabolism and homeostasis, including cholesterol biosynthesis, fatty acid metabolism, and lipid storage, as well as oxidative stress, ER stress and protein quality control, lysosomal function, and intracellular trafficking (Fig. 1D). We next performed high-dimensional weighted gene co-expression network analysis (hdWGCNA)^28^ across symptomatic HD-MSNs to identify coordinated transcriptional programs associated with cluster 2. Among six co-expression modules, M1, M3, and M6 were consistently enriched in cluster 2 relative to clusters 0, 1, and 3 by both module eigengene and UCell-based scoring^29^ (Supplementary Fig. 1B). M1 was associated with lipid metabolism, lipid localization and transport, inflammatory signaling, and oxidative stress; M3 with protein folding and disassembly, autophagy, and mitophagy; and M6 with axonal transport, organelle localization, and microtubule-based movement (Fig. 1E). TOM-based network projection showed that M1, M3, and M6 occupied neighboring regions of the co-expression network (Supplementary Fig. 1C). Representative genes included MGLL, PSAP, and NPC2 in M1; CANX, HSPA8, and BNIP3L in M3; and KIF5C, VPS41, and MAP4 in M6, consistent with the functions of these modules in lipid metabolism, protein folding and autophagy, and intracellular transport. Together, these analyses revealed a coordinated transcriptional state in HD-MSNs derived from symptomatic individuals.

We next asked whether these transcriptional features were also represented in neurons from the human HD brain. We performed pre-ranked gene set enrichment analysis (GSEA) of hdWGCNA modules M1, M3, and M6 using HD-versus-control differential expression signatures from postmortem human caudate direct- and indirect-pathway spiny projection neurons (dSPNs and iSPNs, respectively)^30^. M3 showed significant positive enrichment in both dSPNs and iSPNs, whereas M1 showed significant positive enrichment in dSPNs. In contrast, M6 exhibited significant negative enrichment in dSPNs (Fig. 1F). These findings support the relevance of transcriptional programs related to lipid metabolism, protein folding, and autophagy identified in symptomatic HD-MSNs to neuronal alterations in the human HD caudate. In a complementary analysis, genes enriched in cluster 2 were compared with genes upregulated in HD neurons relative to unaffected controls in two independent postmortem human brain snRNA-seq datasets^5,30^. We identified 20 overlapping genes in striatal D1/D2 MSNs from HD grade 2-4 samples and 23 overlapping genes in caudate DARPP32-positive MSNs from HD grade 3 samples (Supplementary Fig. 1D). These overlapping genes were enriched for processes including phospholipid and sphingolipid catabolism, lysosomal targeting, and responses to misfolded or unfolded proteins, with dataset-specific associations with intracellular iron homeostasis and ferroptosis. Thus, transcriptional signature associated with symptomatic cluster 2 were also represented in neurons from the human HD brain.

### HD-LINC is selectively induced in symptomatic HD-MSNs and enriched in the disease-associated neuronal state

To identify lncRNAs associated with symptomatic HD, we first examined three independent bulk RNA-seq datasets generated from directly reprogrammed MSNs, including two healthy control versus symptomatic HD comparisons and one pre-symptomatic versus symptomatic HD comparison^11,12,31^. In one Ctrl-MSN versus symptomatic HD-MSN dataset, HD-LINC (LINC01139) was the second most highly upregulated transcript overall and the most highly upregulated lncRNA (Fig. 1G). Four lncRNAs were shared across all three comparisons, but HD-LINC exhibited substantially higher expression than the remaining candidates (Supplementary Fig. 1E). IGV visualization further showed robust HD-LINC transcription in symptomatic HD-MSNs but little or no detectable expression in healthy Ctrl-MSNs or pre-HD-MSNs across all three datasets (Supplementary Fig. 1F). RT-qPCR using directly reprogrammed MSNs from age-matched healthy controls (n = 4 donors) and symptomatic HD patients (n = 8 donors) independently confirmed significantly elevated HD-LINC expression in symptomatic HD-MSNs compared with Ctrl-MSNs (Supplementary Fig. 1G).

We next examined HD-LINC expression at single-cell resolution in pre-HD-MSNs and symptomatic HD-MSNs. Notably, HD-LINC was absent or minimally detected in pre-HD-MSNs but selectively induced in symptomatic HD-MSNs, with expression predominantly enriched within cluster 2 (Fig. 1H). TSTD1, a cytoplasmic sulfurtransferase implicated in sulfide metabolism and thioredoxin-dependent redox regulation^32,33^, showed the opposite pattern, with expression readily detected in pre-HD-MSNs but markedly reduced in symptomatic HD-MSNs. In contrast, PLPP2, which encodes a phospholipid phosphatase involved in the metabolism of bioactive lipid phosphates^34^, was increased in symptomatic HD-MSNs. A general neuronal marker SNAP25 was broadly expressed across clusters in both groups, demonstrating the neuronal identity of the analyzed cells (Fig. 1H). Thus, HD-LINC was selectively associated with cluster 2 in the symptomatic HD neuronal state rather than simply reflecting the presence of mutant HTT.

Finally, RNAscope analysis of postmortem human dorsal caudate revealed significantly increased HD-LINC signals in both grade 1 and grade 2-4 HD samples compared with controls, with a further increase in grade 2-4 relative to grade 1 samples (Fig. 1I). Quantification was performed in neuronal regions identified by MAP2 staining, and neuronal localization was further confirmed by phosphorylated neurofilament staining (Supplementary Fig. 2A). RNAscope analysis of directly reprogrammed MSNs similarly showed markedly increased HD-LINC signals in HD-MSNs compared with Ctrl-MSNs (Fig. 1I). Co-analysis with SNAP25 probe further confirmed HD-LINC expression in reprogrammed neurons, with transcripts detected in the nucleus as well as throughout the neuronal soma and neurites (Fig. 1I). Together, these findings identify HD-LINC as a human-specific lincRNA selectively induced within a symptomatic HD neuronal state enriched for transcriptional programs related to lipid metabolism, protein folding, and autophagy, with HD-LINC signals increasing with HD pathological grade in the human caudate.

### HD-LINC contributes to neurodegenerative phenotypes in HD-MSNs

To determine the appropriate time point for phenotypic analyses, we monitored neuronal degeneration during direct reprogramming by measuring Annexin V-positive cells from post-induction day (PID)28 to PID32. Ctrl-MSNs and HD-MSNs showed comparable cell death at PID28 and PID29, whereas HD-MSNs exhibited increased Annexin V-positive cells beginning at PID30, which became more pronounced at PID31 and PID32. We therefore selected PID30 for subsequent phenotypic analyses (Supplementary Fig. 3A). We next investigated whether reducing HD-LINC expression could alleviate neurodegenerative phenotypes in HD-MSNs. HD-LINC knockdown (HD-shLINC) significantly decreased Annexin V-positive cells compared with control shRNA (HD-shCon), whereas knockdown did not affect Annexin V-positive cells in Ctrl-MSNs (Fig. 2A, B), indicating that the protective effect of HD-LINC knockdown was preferentially observed in the HD disease context. We next examined dystrophic neurites, identified by bead- or bulge-like structures along TUBB3-positive neurites. HD-LINC knockdown significantly reduced dystrophic neuritic events per 100 μm of neurite length (Fig. 2C), which was independently confirmed by quantifying the percentage of neurites exhibiting dystrophic morphology (Supplementary Fig. 3B). HD-LINC knockdown also reduced the DNA damage marker 53BP1 (Fig. 2D) and markedly decreased the percentage of neurons containing mutant HTT (mHTT) inclusion bodies (Fig. 2E). Thus, HD-LINC repression attenuated multiple HD-associated phenotypes, including neuronal death, neuritic degeneration, DNA damage, and mHTT aggregation. To determine whether increased HD-LINC expression could instead exacerbate these phenotypes, we overexpressed HD-LINC in HD-MSNs. HD-LINC overexpression increased Annexin V-positive cells, dystrophic neuritic events, 53BP1-positive area, and the percentage of neurons containing mHTT inclusion bodies compared with LacZ control (Fig. 2F-I). Increased neuritic degeneration was independently confirmed by quantifying the percentage of neurites exhibiting dystrophic morphology (Supplementary Fig. 3B). Together, these loss- and gain-of-function experiments demonstrate that HD-LINC contributes to neuronal death, neuritic degeneration, DNA damage, and mutant HTT aggregation in patient-derived HD-MSNs.

**Figure 2.**
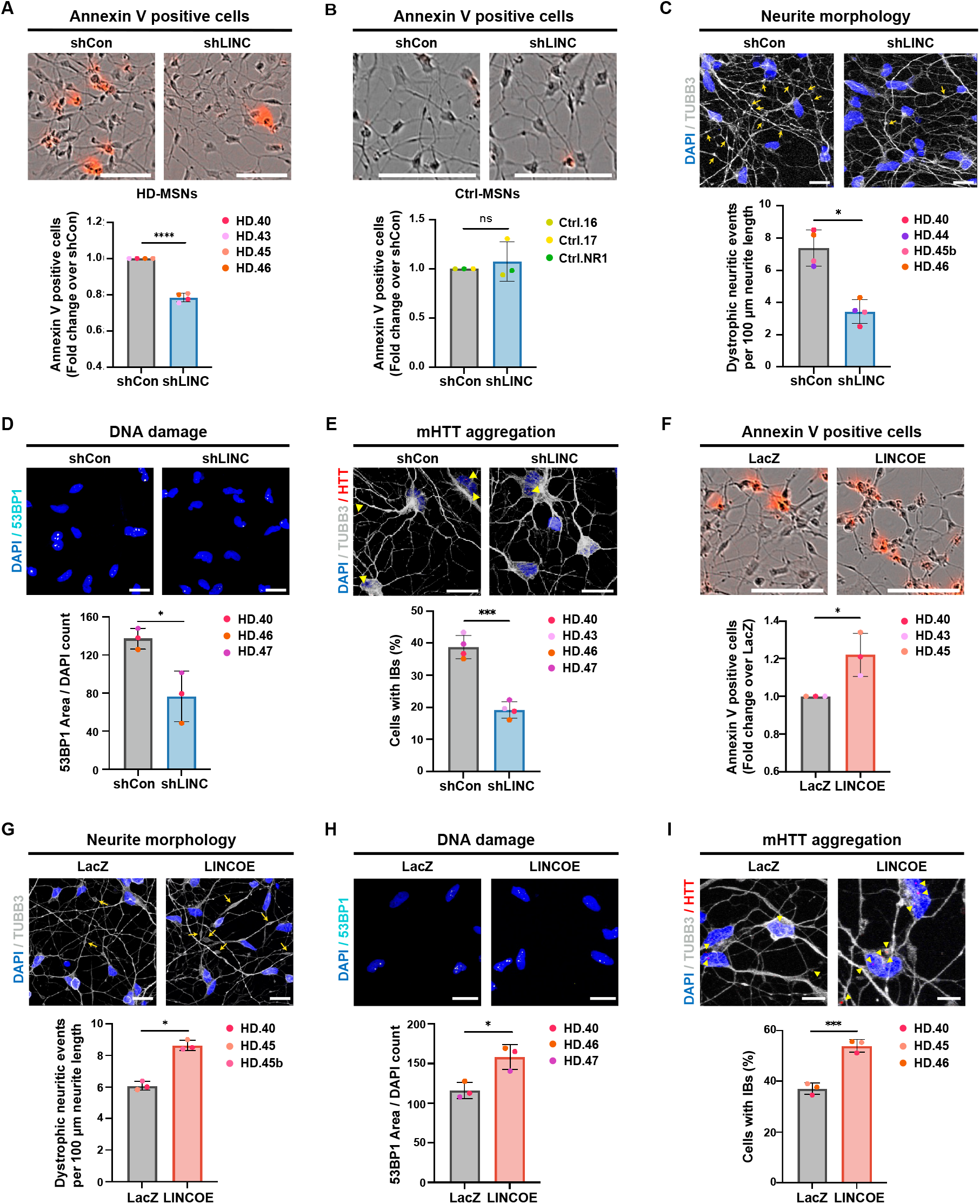
HD-LINC repression alleviates HD-associated neurodegenerative phenotypes in HD-MSNs. **(A)** Representative images (top) and quantification (bottom) of Annexin V-positive cells in HD-MSNs expressing control shRNA (shCon) or shRNA targeting HD-LINC (shLINC). **(B)** Representative images (top) and quantification (bottom) of Annexin V-positive cells in Ctrl-MSNs expressing shCon or shLINC. **(C–E)** Representative images (top) and quantification (bottom) of dystrophic neurite morphology **(C)**, DNA damage, quantified as 53BP1-positive area per DAPI-positive cell **(D)**, and mutant HTT (mHTT) inclusion bodies (IBs) **(E)** in HD-MSNs expressing shCon or shLINC. (F–I) Representative images (top) and quantification (bottom) of Annexin V-positive cells **(F)**, dystrophic neurite morphology **(G)**, DNA damage, quantified as 53BP1-positive area per DAPI-positive cell **(H)**, and mHTT IBs **(I)** in HD-MSNs expressing LacZ or an HD-LINC overexpression construct (LINC OE). Dystrophic neurites were quantified as beaded or bulged neuritic events per 100 μm of neurite length. Each dot represents one independent reprogrammed MSN line. Sample sizes were n = 3 for B, D, and F–I and n = 4 for A, C, and E. Statistical significance was determined by two-tailed paired t-test. Data are mean ± s.e.m. ns, not significant; *P < 0.05, ***P < 0.001, ****P < 0.0001. Scale bars, 20 μm (except Fig. 2I) and 10 μm (Fig. 2I).

### HD-LINC regulates transcriptional programs associated with lipid metabolism and protein clearance

To investigate transcriptional changes following HD-LINC repression, we performed bulk RNA-seq and targeted lipidomics in symptomatic HD-MSNs expressing control shRNA (HD-shCon) or shRNA targeting HD-LINC (HD-shLINC) (Fig. 3A). As expected, differential gene expression analysis identified HD-LINC (LINC01139) as the most significantly downregulated transcript following its knockdown (Fig. 3B). Among the prominently altered genes, LPL and PTGS1, involved in lipid metabolism and signaling, were downregulated, whereas PRELID1, involved in mitochondrial phospholipid transport^35^, was increased. SORL1, a regulator of endosomal and lysosomal trafficking^36^, and HSPA6, a stress-inducible HSP70-family protein associated with protein disaggregation and refolding machinery^37^, were also increased (Fig. 3B), linking HD-LINC repression to changes in lipid biology, intracellular trafficking, and protein homeostasis. Gene ontology (GO) analysis of genes upregulated following HD-LINC knockdown revealed enrichment of protein degradation and ER-associated degradation pathways, together with phospholipid transport, lipoprotein metabolism, triglyceride catabolism, and lysosomal and intracellular trafficking (Fig. 3C). Genes downregulated following HD-LINC repression were enriched for pathways involving vesicle fusion and exocytosis, synaptic organization, actin regulation, oxidative stress, and triglyceride-rich lipoprotein remodeling (Supplementary Fig. 4A). These analyses identified lipid metabolism, protein clearance, and intracellular trafficking as major transcriptional processes altered by HD-LINC repression. We next performed weighted gene co-expression network analysis (WGCNA)^38^ to identify coordinated transcriptional programs associated with HD-LINC knockdown. Six co-expression modules were significantly associated with the HD-shLINC condition (Fig. 3D). Among the positively associated modules, the darkgreen module was enriched for cholesterol and lipid biosynthesis and fatty acid metabolism, whereas the purple module was enriched for lysosomal targeting, proteasomal protein degradation, and vesicle-mediated transport (Fig. 3E). Other associated modules included processes related to neuronal morphology, oxidative and iron homeostasis, lysosomal localization, and proteasomal degradation (Supplementary Fig. 4B). Network analysis further resolved interconnected lipid biosynthetic and fatty acid metabolic subnetworks within the darkgreen module, including HMGCR, INSIG1, and ACBD5, and lysosomal/vesicular trafficking and proteasomal subnetworks within the purple module, including SORL1, SORT1, and SKP1 (Fig. 3F). Thus, transcriptional changes following HD-LINC repression were organized within coordinated lipid metabolic and protein clearance networks.

**Figure 3.**
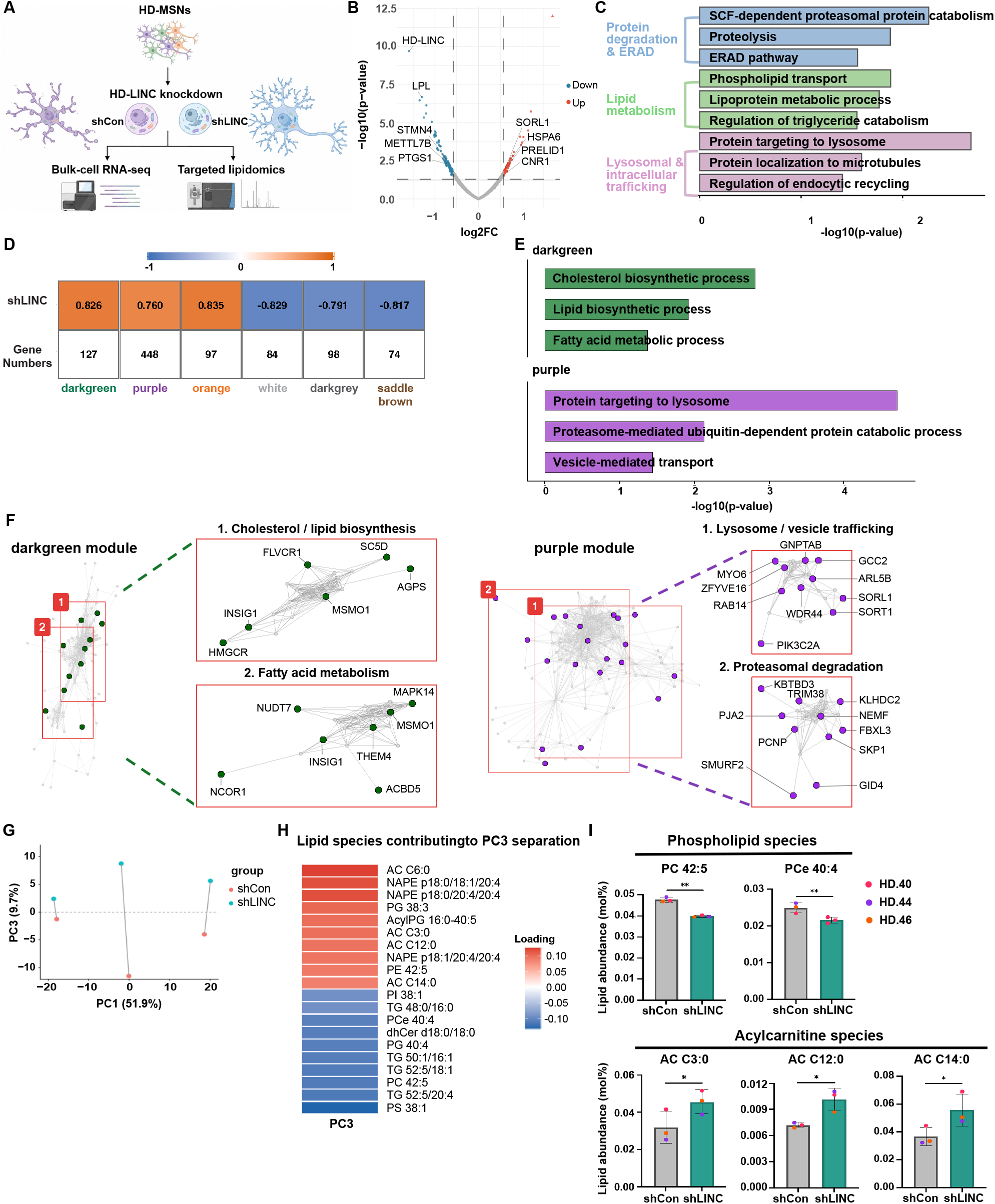
HD-LINC knockdown alters lipid metabolic and protein degradation pathways and remodels the HD-MSN lipidome. **(A)** Schematic of HD-LINC knockdown followed by bulk-cell RNA-seq and targeted lipidomic analyses in HD-MSNs expressing control shRNA (shCon) or shRNA targeting HD-LINC (shLINC). **(B)** Volcano plot of differentially expressed genes between HD-shCon and HD-shLINC MSNs. Blue, downregulated genes; red, upregulated genes. **(C)** Gene ontology (GO) terms associated with genes upregulated following HD-LINC knockdown, highlighting protein degradation and ER-associated degradation (ERAD), lipid metabolism, and lysosomal and intracellular trafficking pathways. Bars represent −log_10_ (P value). **(D)** WGCNA module–trait correlations between gene co-expression modules identified from bulk RNA-seq and HD-LINC knockdown. Correlation coefficients and the number of genes contained within each module are shown. **(E)** GO terms associated with the darkgreen and purple WGCNA modules, highlighting lipid biosynthetic and metabolic processes and pathways related to lysosomal targeting, proteasomal degradation, and vesicle-mediated transport. Bars represent −log_10_ (P value). **(F)** Co-expression networks of the darkgreen and purple WGCNA modules. Selected subnetworks associated with cholesterol/lipid biosynthesis, fatty acid metabolism, lysosome/vesicle trafficking, and proteasomal degradation are shown at higher magnification. **(G)** Principal component analysis of targeted lipidomic profiles from HD-shCon and HD-shLINC MSNs showing PC1 and PC3. Samples derived from the same independent HD-MSN line are connected by lines. **(H)** Lipid species contributing to PC3 separation between HD-shCon and HD-shLINC MSNs. Bars indicate PC3 loadings. **(I)** Quantification of selected significantly altered phospholipid and acylcarnitine species in HD-shCon and HD-shLINC MSNs, including PC 42:5, PCe 40:4, AC C3:0, AC C12:0, and AC C14:0. Bulk-cell RNA-seq and WGCNA analyses were performed using n = 4 independent HD-MSN lines per condition. Targeted lipidomic analyses were performed using n = 3 independent HD-MSN lines per condition. For quantifications in I, each dot represents one independent HD-MSN line. Statistical significance in I was determined by two-tailed paired t-test. Data are mean ± s.e.m. *P < 0.05, **P < 0.01.

### HD-LINC knockdown remodels lipid profiles in HD-MSNs

Because transcriptomic analyses consistently implicated lipid-associated processes, we next performed lipidomic profiling of paired HD-shCon and HD-shLINC samples from three independent HD-MSN lines (Fig. 3A). Principal component analysis (PCA) revealed substantial donor-specific variation along PC1, accounting for 61.9% of the total variance, whereas paired HD-shCon and HD-shLINC samples showed a consistent shift along PC3, accounting for 17.7% of the variance (Fig. 3G). Paired PC3 scores increased following HD-LINC knockdown across all three HD-MSN lines (Supplementary Fig. 4C), indicating a reproducible lipidomic shift despite donor-specific variation. Analysis of PC3 loadings showed that acylcarnitine (AC) species contributed positively to the HD-shLINC-associated shift, whereas phosphatidylcholine (PC), ether phosphatidylcholine (PCe), and several triglyceride (TG) species, which are enriched in LDs, contributed in the opposite direction (Fig. 3H). Several lipid species rich in polyunsaturated fatty acids (PUFAs) were correlated with HD-LINC levels. At the individual species level, PC 42:5 and PCe 40:4 were significantly decreased following HD-LINC knockdown, whereas AC C3:0, AC C12:0, and AC C14:0 were increased (Fig. 3I). Lipid-class analysis further showed increased total acylcarnitines and decreased monoacylglycerols in shHD-LINC relative to HD-shCon (Supplementary Fig. 4D). Because PUFA phospholipids are susceptible to lipid peroxidation^39^, acylcarnitines are intermediates in mitochondrial fatty acid transport and oxidation^40^, and triglycerides are major neutral lipid components of lipid droplets^41^, these lipidomic changes suggest that HD-LINC may influence the propensity of lipid to peroxidation, lipid metabolism, and LD biology.

### HD-LINC promotes dysregulated neuritic lipid droplet accumulation associated with reduced LD mobility in HD-MSNs

We next examined lipid droplet (LD) dynamics in HD-MSNs. Live-cell imaging of BODIPY-labeled LDs directly visualized movement of LDs from the soma into neurites, revealing dynamic distribution of LDs between neuronal compartments. Importantly, when we measured the mobility of LDs along neurites, LD mobilities significantly differed according to LD sizes. Large neuritic LDs showed significantly lower net displacement than small LDs over the 5-min imaging period, and enlarged LDs were frequently retained within neurites with limited movement (Fig. 4A). These findings indicate an association between reduced mobility of enlarged LDs and their retention within neurites, providing a potential dynamic basis for neuritic LD accumulation. Representative high-magnification time-lapse imaging is shown in Supplementary Movies 1 and 2.

**Figure 4.**
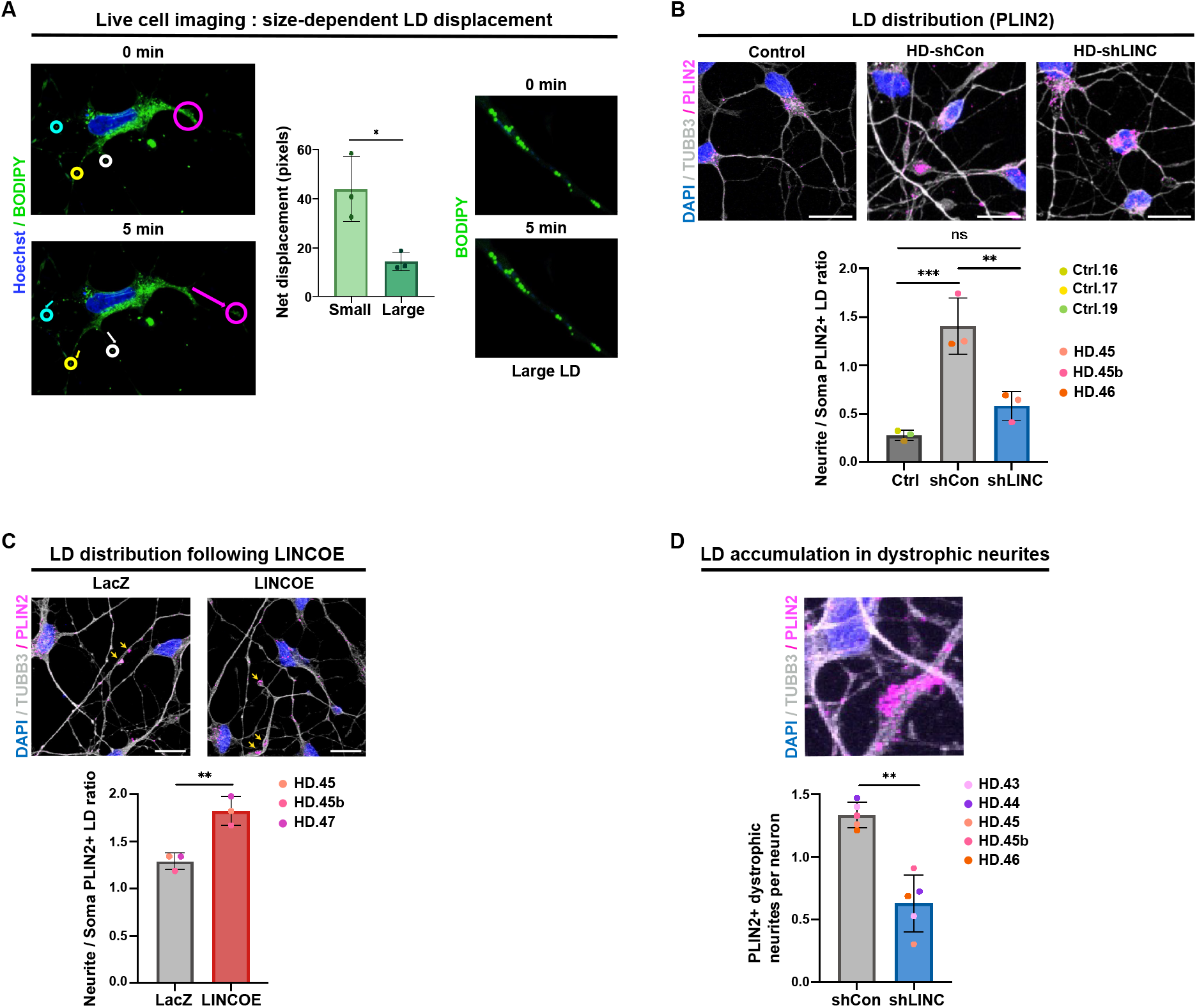
HD-LINC knockdown reduces neuritic lipid droplet accumulation in HD-MSNs. **(A)** Live-cell imaging of BODIPY-labeled lipid droplets (LDs) in HD-MSNs showing size-dependent neuritic LD displacement. Representative time-lapse images at 0 and 5 min show movement of individual LDs, indicated by colored circles. Quantification shows net displacement of small and large neuritic LDs over 5 min. Representative high-magnification time-lapse images show limited movement of a large neuritic LD. **(B)** Representative images (top) and quantification (bottom) of the neurite-to-soma PLIN2-positive LD ratio in Ctrl-MSNs, HD-MSNs expressing control shRNA (HD-shCon), and HD-MSNs expressing shRNA targeting HD-LINC (HD-shLINC). **(C)** Representative images (top) and quantification (bottom) of the neurite-to-soma PLIN2-positive LD ratio in HD-MSNs expressing LacZ or an HD-LINC overexpression construct (LINC OE). **(D)** Representative image (top) and quantification (bottom) of PLIN2-positive beaded or bulged dystrophic neurites per neuron in HD-shCon and HD-shLINC MSNs. For live-cell imaging in A, one independent HD-MSN line was used; each dot represents one neuron, and each neuron-level value represents the mean net displacement of five small or five large neuritic LDs. For B, n = 3 Ctrl-MSN lines and n = 3 HD-MSN lines were analyzed. For C, n = 3 independent HD-MSN lines per condition were analyzed. For D, n = 5 independent HD-MSN lines per condition were analyzed. Each dot in B–D represents one independent reprogrammed MSN line. Statistical significance was determined by two-tailed paired t-test for LD displacement in A, one-way ANOVA followed by Tukey’s multiple-comparisons test in B, and two-tailed paired t-test in C and D. Data are mean ± s.e.m. ns, not significant; *P < 0.05, **P < 0.01, ***P < 0.001. Scale bars, 20 μm.

Given the association between LD size, reduced mobility, and neuritic retention, we next examined whether HD-LINC regulates LD accumulation and spatial distribution in HD-MSNs. Importantly, immunostaining for PLIN2 showed that Ctrl-MSNs displayed predominantly soma-localized LDs, whereas HD-shCon MSNs exhibited prominent neuritic LD accumulation. Importantly, HD-LINC knockdown markedly reduced neuritic LD accumulation and restored a control-like soma-enriched LD distribution, as quantified by a significant decrease in the neurite-to-soma PLIN2-positive LD ratio (Fig. 4B). Independent staining with the neutral lipid dye BODIPY 493/503^42^ similarly confirmed reduced neuritic LD accumulation following HD-LINC knockdown and a decreased neurite-to-soma LD ratio (Supplementary Fig. 5A). Conversely, overexpressing HD-LINC increased neuritic LD accumulation, reflected by an increased neurite-to-soma PLIN2-positive LD ratio compared with LacZ-expressing HD-MSNs (Fig. 4C). Together, these reduction- and gain-of-function experiments demonstrate that HD-LINC promotes dysregulated neuritic LD accumulation and regulates LD spatial distribution in HD-MSNs. We next examined whether neuritic LD accumulation was associated with dystrophic neurites. PLIN2-positive LDs were frequently observed within dystrophic neurites, and HD-LINC knockdown significantly reduced the number of PLIN2-positive dystrophic neurites per neuron (Fig. 4D). Consistent with the increased somatic localization of LDs following HD-LINC knockdown, we also observed an increased association of mHTT aggregates with PLIN2-positive LDs, predominantly within the soma (Supplementary Fig. 5B). Together, these findings identify dysregulated neuritic LD accumulation as a major phenotype regulated by HD-LINC and link altered LD spatial organization and reduced LD mobility to neuritic vulnerability in HD-MSNs.

### HD-LINC promotes oxidative lipid stress linked to aberrant LDs and mutant HTT aggregation in HD-MSNs

We next asked whether LD accumulation was associated with oxidative lipid damage within dystrophic neurites. Co-immunostaining for PLIN2, a lipid droplet surface protein^43^, and 4-HNE, a reactive aldehyde generated during lipid peroxidation^44^, revealed PLIN2-positive/4-HNE-positive dystrophic neurites in HD-shCon MSNs, demonstrating that LD accumulation and lipid peroxidation co-occur within the same dystrophic neurites (Fig. 5A). HD-LINC knockdown significantly reduced the number of these double-positive dystrophic neurites, indicating reduced co-occurrence of LD accumulation and oxidative lipid damage within dystrophic neurites.

**Figure 5.**
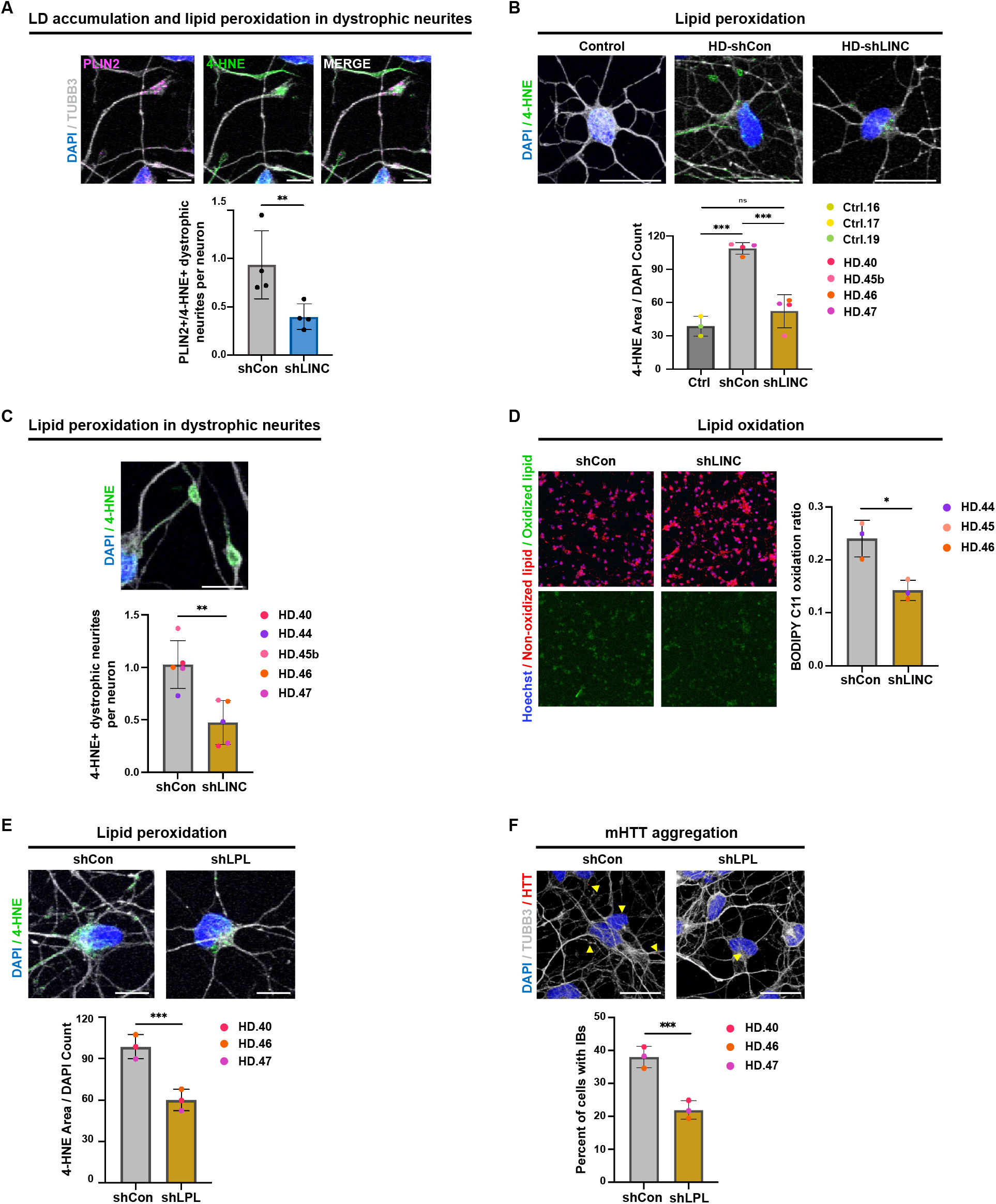
HD-LINC knockdown reduces lipid peroxidation in HD-MSNs. **(A)** Representative images (top) and quantification (bottom) of PLIN2- and 4-hydroxynonenal (4-HNE)-positive beaded or bulged dystrophic neurites in HD-MSNs expressing control shRNA (HD-shCon) or shRNA targeting HD-LINC (HD-shLINC). **(B)** Representative images (top) and quantification (bottom) of 4-HNE-positive area per DAPI-positive cell in Ctrl-MSNs, HD-shCon MSNs, and HD-shLINC MSNs. **(C)** Representative image (top) and quantification (bottom) of 4-HNE-positive beaded or bulged dystrophic neurites per neuron in HD-shCon and HD-shLINC MSNs. **(D)** Live-cell imaging of lipid oxidation using BODIPY 581/591 C11 in HD-shCon and HD-shLINC MSNs. Left, non-oxidized and oxidized BODIPY C11 signals; right, quantification of the BODIPY C11 oxidation ratio. **(E)** Representative images (top) and quantification (bottom) of 4-HNE-positive area per DAPI-positive cell in HD-MSNs expressing shCon or shRNA targeting LPL (shLPL). **(F)** Representative images (top) and quantification (bottom) of mutant HTT (mHTT) inclusion bodies (IBs) in HD-MSNs expressing shCon or shLPL. Quantification of PLIN2/4-HNE-positive dystrophic neurites in A was performed using one independent HD-MSN line with four wells per condition; individual dots represent wells. For B, n = 3 Ctrl-MSN lines and n = 4 HD-MSN lines were analyzed. For C, n = 5 independent HD-MSN lines per condition were analyzed. For D–F, n = 3 independent HD-MSN lines per condition were analyzed. Each dot in B–F represents one independent reprogrammed MSN line. Statistical significance was determined by two-tailed unpaired t-test in A, one-way ANOVA followed by Tukey’s multiple-comparisons test in B, and two-tailed paired t-test in C–F. Data are mean ± s.e.m. ns, not significant; *P < 0.05, **P < 0.01, ***P < 0.001. Scale bars, 10 μm (Fig. 5A, C, and E) and 20 μm (Fig. 5B and F).

Given the observed co-occurrence of LD accumulation and lipid peroxidation within dystrophic neurites, we next investigated whether HD-LINC more broadly regulates oxidative lipid stress, a necessary step for lipid-peroxidation-mediated cell death, in HD-MSNs. We first examined lipid peroxidation by immunostaining for 4-HNE. HD-shCon MSNs showed markedly increased 4-HNE accumulation compared with Ctrl-MSNs, and HD-LINC knockdown significantly reduced 4-HNE levels relative to HD-shCon (Fig. 5B). We next examined lipid peroxidation in dystrophic neurites. HD-LINC knockdown significantly reduced the number of 4-HNE-positive dystrophic neurites per neuron compared with HD-shCon (Fig. 5C), indicating reduced oxidative lipid damage associated with dystrophic neurites following HD-LINC repression. To independently assess lipid oxidation, we measured oxidation of the live-cell lipid peroxidation reporter C11-BODIPY^45^. HD-LINC knockdown significantly reduced the C11-BODIPY oxidation ratio compared with HD-shCon (Fig. 5D), further demonstrating that HD-LINC repression reduces oxidative lipid stress in HD-MSNs.

Our RNA-seq analysis identified LPL as one of the lipid metabolic genes most strongly downregulated following HD-LINC knockdown (Fig. 3B). Given its reported role in neuronal lipid uptake and storage^46,47^, we next tested whether the effects of HD-LINC on lipid peroxidation in HD-MSNs are mediated solely through LPL. We first confirmed efficient HD-LINC knockdown together with a marked reduction in LPL expression in HD-shLINC MSNs (Supplementary Fig. 6A). We found that independent LPL knockdown decreased 4-HNE accumulation (Fig. 5E) and the percentage of neurons containing mHTT inclusions (Fig. 5F) compared with HD-shCon. To further investigate whether lipid peroxidation contributes to mHTT aggregation, we treated HD-MSNs with ferrostatin-1 (Fer-1), a lipid radical-trapping antioxidant^48^. Fer-1 treatment reduced mHTT aggregates across multiple patient-derived HD-MSN lines (Supplementary Fig. 6B). However, in contrast to HD-LINC repression, LPL knockdown did not significantly alter the neurite-to-soma PLIN2-positive LD ratio in HD-MSNs (Supplementary Fig. 6C), suggesting that LPL reduction alone was insufficient to reproduce the effect of HD-LINC repression on LD spatial distribution. While it remains unclear how HD-LINC contributes to the dysregulation of LD trafficking, our results overall demonstrate the multiple functionality of HD-LINC in promoting oxidative lipid stress, mHTT inclusion burden, and aberrant neuritic LD accumulation.

### Antisense oligonucleotide-mediated repression of HD-LINC recapitulates the protective effects of shRNA-mediated knockdown

Because shRNA-mediated repression of HD-LINC attenuated multiple HD-associated phenotypes, we next asked whether these effects could be reproduced using an antisense oligonucleotide (ASO) targeting HD-LINC. We first evaluated ASO-mediated HD-LINC repression across increasing concentrations. HD-LINC ASO treatment produced a dose-dependent reduction in HD-LINC expression, with 7.5 nM reducing HD-LINC levels by approximately 80% relative to the non-targeting control ASO (Fig. 6A). Consistent with the transcriptional changes observed following shRNA-mediated HD-LINC knockdown, LPL expression closely tracked with HD-LINC expression across ASO doses (Supplementary Fig. 7A). We next examined whether ASO-mediated HD-LINC repression reduced oxidative lipid stress. Treatment with 7.5 nM HD-LINC ASO significantly decreased cellular 4-HNE levels compared with the control ASO (Fig. 6B). HD-LINC ASO treatment also significantly reduced the number of 4-HNE-positive dystrophic neurites per neuron (Fig. 6C), indicating reduced oxidative lipid damage associated with dystrophic neurites. We then asked whether ASO-mediated HD-LINC repression also affected neuritic LD accumulation. HD-LINC ASO treatment significantly reduced the neurite-to-soma PLIN2-positive LD ratio, consistent with reduced neuritic LD accumulation and a more soma-enriched LD distribution (Fig. 6D). In addition, ASO treatment significantly decreased the number of PLIN2-positive dystrophic neurites per neuron (Fig. 6E). Thus, ASO-mediated repression of HD-LINC recapitulated the effects of shRNA-mediated knockdown on both oxidative lipid stress and dysregulated neuritic LD accumulation. Finally, we examined whether these improvements in lipid dysregulation were accompanied by reductions in neurodegenerative phenotypes. HD-LINC ASO treatment significantly decreased the percentage of neurons containing mutant HTT inclusion bodies (Fig. 6F) and reduced Annexin V-positive cells compared with the control ASO (Fig. 6G). Importantly, ASO-mediated repression of HD-LINC recapitulated the protective effects of shRNA-mediated knockdown by reducing oxidative lipid stress, dysregulated neuritic LD accumulation, mutant HTT aggregation, and neuronal death in HD-MSNs.

**Figure 6.**
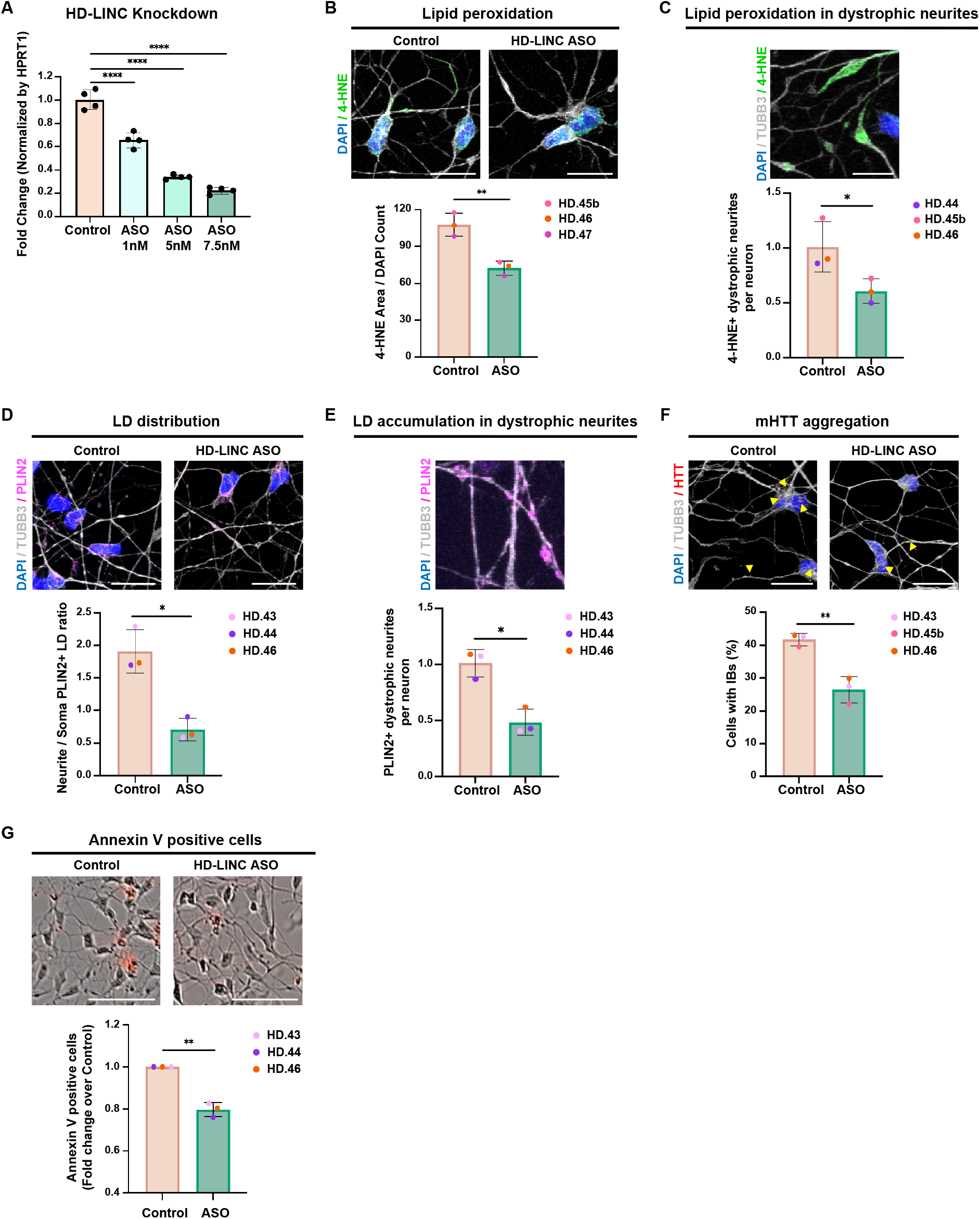
Antisense oligonucleotide-mediated repression of HD-LINC recapitulates the protective effects of HD-LINC knockdown in HD-MSNs. **(A)** RT–qPCR analysis of HD-LINC expression in HD-MSNs treated with increasing concentrations of an antisense oligonucleotide (ASO) targeting HD-LINC. **(B)** Representative images (top) and quantification (bottom) of 4-hydroxynonenal (4-HNE)-positive area per DAPI-positive cell in HD-MSNs treated with control ASO or HD-LINC ASO. **(C)** Representative image (top) and quantification (bottom) of 4-HNE-positive beaded or bulged dystrophic neurites per neuron in HD-MSNs treated with control ASO or HD-LINC ASO. **(D)** Representative images (top) and quantification (bottom) of the neurite-to-soma PLIN2-positive lipid droplet (LD) ratio in HD-MSNs treated with control ASO or HD-LINC ASO. **(E)** Representative image (top) and quantification (bottom) of PLIN2-positive beaded or bulged dystrophic neurites per neuron in HD-MSNs treated with control ASO or HD-LINC ASO. **(F)** Representative images (top) and quantification (bottom) of mutant HTT (mHTT) inclusion bodies (IBs) in HD-MSNs treated with control ASO or HD-LINC ASO. **(G)** Representative images (top) and quantification (bottom) of Annexin V-positive cells in HD-MSNs treated with control ASO or HD-LINC ASO. For dose-response analyses in A, one independent HD-MSN line was used, with four wells analyzed per ASO concentration; each dot represents one well. For quantifications in B–G, n = 3 independent HD-MSN lines per condition were analyzed, and each dot represents one independent HD-MSN line. Statistical significance was determined by one-way ANOVA followed by Dunnett’s multiple-comparisons test in A and two-tailed paired t-test in B–G. Data are mean ± s.e.m. *P < 0.05, **P < 0.01, ****P < 0.0001. Scale bars, 20 μm.

## Discussion

Findings in this study identify HD-LINC as a human-specific long noncoding RNA associated with symptomatic HD and neuronal vulnerability. HD-LINC was selectively induced in symptomatic HD-MSNs and enriched in a neuronal state characterized by changes in lipid metabolism, protein quality control, autophagy, and intracellular trafficking. Functional perturbation further demonstrated that HD-LINC promotes dysregulated neuritic lipid droplet (LD) accumulation and increased lipid peroxidation, whereas its repression restored a more control-like LD distribution, reduced lipid peroxidation, and mitigated mutant HTT aggregation and neuronal degeneration. These findings reveal a previously unrecognized human-specific regulatory mechanism linking altered lipid metabolism to vulnerability in HD-MSNs. An important feature of HD-LINC is its selective association with the symptomatic HD state. Despite the presence of mutant HTT before clinical onset, HD-LINC was minimally detected in pre-symptomatic HD-MSNs but strongly induced in symptomatic HD-MSNs. Related transcriptional signatures in independent postmortem HD neuronal datasets^5,30^ further support the relevance of disease-stage-specific molecular programs that emerge beyond the presence of the causative mutation and may contribute to neuronal vulnerability.

Previous studies characterized LINC01139 as LINK-A in breast cancer cells, where it was identified as a lipid-binding lncRNA that interacts with phosphatidylinositol-3,4,5-trisphosphate (PIP3) and facilitates AKT activation^21^. However, its function in human neurons has remained unknown. Our findings reveal a role for HD-LINC in neuronal lipid regulation and neurodegeneration in human HD-MSNs. Transcriptomic and lipidomic analyses identified coordinated changes in lipid metabolism, protein degradation, and lysosomal and intracellular trafficking, together with altered phospholipid and acylcarnitine profiles, supporting a broader role for HD-LINC in neuronal lipid homeostasis. It remains to be determined whether the lipid-binding properties previously described for LINC01139 are retained in neurons and which molecular interactions mediate the effects of HD-LINC on neuronal lipid biology.

LD accumulation has been reported in HD and other neurodegenerative disorders^16,17,49^, but these observations have largely described the presence of LDs without defining their intracellular organization or functional significance. Our findings identify the subcellular distribution of LDs as an additional dimension of neuronal lipid dysregulation in HD. HD-MSNs displayed prominent neuritic LD accumulation, whereas HD-LINC repression restored a more soma-enriched distribution and HD-LINC overexpression further enhanced neuritic LD accumulation. Larger neuritic LDs also exhibited reduced net displacement. These observations suggest that LD mobility and compartmental redistribution may be important features of how lipids are spatially organized within neurons. Given the highly polarized architecture of neurons and their dependence on long-range intracellular transport^50^, the consequences of LD accumulation may depend not only on the amount of stored lipid but also on where LDs are positioned within the cell. The enrichment of genes involved in intracellular trafficking, organelle localization, and microtubule-based movement in the symptomatic HD neuronal state further raises the possibility that changes in neuronal transport processes may contribute to the altered spatial organization of LDs. Whether changes in LD positioning and mobility alter local trafficking, organelle interactions, or lipid utilization will be important to determine in future studies. The spatial association of LD accumulation with dystrophic neurites, together with the co-occurrence of LDs and oxidative lipid damage within these degenerating regions, further links altered LD organization to neuritic vulnerability in HD-MSNs. Together, these findings suggest that HD-LINC-dependent neuritic LD accumulation represents more than a generalized increase in cellular lipid storage, identifying altered LD spatial organization as a distinct and previously underexplored feature of HD patient-derived neurons. How HD-LINC contributes to altered LD trafficking and retention within neurites, however, remains to be determined. Elucidating these mechanisms and defining the functional consequences of altered LD positioning will be important for understanding how neuronal lipid homeostasis becomes disrupted in HD.

Oxidative lipid damage, a key feature of ferroptosis, has been implicated in HD^51–53^, but the RNA-dependent regulatory mechanisms contributing to lipid peroxidation remain poorly understood. Our findings identify the disease-induced, human-specific lncRNA HD-LINC as a previously unrecognized regulator of oxidative lipid damage in human HD-MSNs. In addition, LPL knockdown reduced lipid peroxidation and mutant HTT inclusion burden without restoring LD spatial distribution, in contrast to HD-LINC repression. Although an association between oxidative lipid damage and mHTT aggregation has previously been reported in HD mouse models using the less specific antioxidant NDGA^54^, the convergent effects of HD-LINC repression, LPL knockdown, and Fer-1 treatment provide complementary genetic and pharmacological evidence supporting a relationship between oxidative lipid stress and mHTT inclusions in human patient-derived HD-MSNs. Although LPL is known to regulate neuronal lipid uptake and storage^46,47^, how these functions relate to oxidative lipid damage in HD-MSNs remains unclear. These findings suggest that HD-LINC regulates oxidative lipid damage and LD spatial organization through at least partially distinct downstream mechanisms that cannot be fully explained by LPL alone.

Whereas current HD therapeutic strategies increasingly target mutant HTT itself^55,56^, our findings suggest that disease-induced lncRNAs may provide an additional means of targeting downstream mechanisms of neuronal vulnerability. ASO-mediated repression of HD-LINC recapitulated the major protective effects of genetic knockdown, providing proof of principle that a human-specific noncoding RNA can be pharmacologically targeted to modify disease-associated phenotypes. Future studies will be required to determine the durability, safety, and therapeutic potential of HD-LINC repression in additional human models and more complex experimental systems.

The human-specific nature of HD-LINC also highlights the importance of human neuronal models for identifying mechanisms that cannot be readily studied in conventional animal systems. Direct neuronal reprogramming preserves donor-associated aging and disease-related states that may be important for capturing symptomatic-stage mechanisms, whereas pluripotent reprogramming may attenuate such features^9,10,12^. The absence of an endogenous HD-LINC ortholog in rodents, however, presents a challenge for defining its physiological function and therapeutic potential in vivo. Future studies should define the molecular interactions through which HD-LINC regulates neuritic LD accumulation and determine how altered LD spatial organization relates to neuronal vulnerability during disease progression. More broadly, our findings suggest that human-specific lncRNAs may represent an underexplored regulatory layer linking disease-state-associated molecular changes to neuronal lipid dysregulation and selective vulnerability in HD and potentially other neurodegenerative disorders.

## METHODS

### Plasmids, shRNAs, antisense oligonucleotides, and cell lines

HD-LINC refers to LINC01139. The construction of plasmids used for MSN reprogramming has been previously described^8,11,12,57^ and the corresponding vectors are publicly available from Addgene, including pTight-9/9*–124-BclxL, rtTA-N144, pmCTIP2-N106, phMYT1L-N174, phDLX1-N174, and phDLX2-N174. Lentiviral shRNAs targeting HD-LINC or LPL and control shRNA were used for loss-of-function experiments. For HD-LINC overexpression, the full-length HD-LINC sequence was cloned and ligated into the N174 lentiviral vector, and a LacZ-expressing vector was used as the control. Affinity Plus phosphorothioate (AP-PS) antisense oligonucleotides (ASOs) targeting HD-LINC and control ASO were obtained from IDT. Adult dermal fibroblasts from symptomatic HD patients, pre-symptomatic HD mutation carriers, and unaffected healthy control individuals were obtained from the Coriell Institute for Medical Research. Donor information, including sex, age at biopsy, CAG repeat length, and experiments performed with each line, is provided in Supplementary Table 1.

### Primary cell culture

Adult human fibroblasts were cultured in fibroblast medium composed of DMEM (high glucose and no glutamine) supplemented with 15% FBS, 0.01% β-mercaptoethanol, 1% nonessential amino acids, 1% sodium pyruvate, 1% GlutaMAX, 1% 1 M HEPES buffer solution, and 1% penicillin/streptomycin solution. Cells were maintained for up to 15 passages.

### Lentivirus preparation

Lentiviral production was performed as previously described^12,57^. Briefly, Lenti-X 293T cells were transfected with individual lentiviral vectors together with the packaging plasmids psPAX2 and pMD2.G using polyethyleneimine. Supernatants were collected 72 h after transfection, filtered through 0.45-μm PES membranes, and concentrated using Lenti-X Concentrator. Concentrated lentiviruses were further purified by centrifugation through a 20% sucrose cushion at 70,000g for 2 h at 4 °C. Viral pellets were resuspended in 10% sucrose solution and stored at −80 °C until use.

### MicroRNA-mediated direct reprogramming of human MSNs

Direct neuronal reprogramming of human fibroblasts to MSNs was performed as previously described^8,11,12,57^. Briefly, human fibroblasts were transduced with a lentiviral cocktail containing pTight-9/9*–124-BclxL, rtTA, CTIP2, DLX1, DLX2, and MYT1L. One day after transduction (PID1), cells were washed and maintained in fibroblast medium containing doxycycline (DOX; 1 μg ml−1). Puromycin (3 μg ml−1) was added at PID3, and cells were replated onto poly-ornithine-, laminin-, and fibronectin-coated coverslips or culture plates at PID5. At PID6, medium was changed to Neurobasal-A containing B-27 Plus, GlutaMAX, DOX (1 μg ml−1), valproic acid (1 mM), dibutyryl cAMP (200 μM), BDNF (10 ng ml−1), NT-3 (10 ng ml−1), retinoic acid (1 μM), RevitaCell Supplement (1×), ascorbic acid (200 μM), puromycin (3 μg ml−1), blasticidin (3 μg ml−1), and geneticin (300 μg ml−1). Blasticidin and geneticin were discontinued after PID10. Beginning at PID14, BrainPhys medium containing NeuroCult SM1 neuronal supplement and N2 Supplement-A was introduced by half-volume medium replacement. DOX was replenished every 2 d and half-volume medium changes were performed every 4 d. RevitaCell and ascorbic acid were discontinued after PID21, and puromycin was maintained until the indicated experimental endpoint.

### Gene perturbation and pharmacological treatments

For shRNA-mediated knockdown experiments, directly reprogrammed MSNs were transduced with lentiviruses expressing shCon, shLINC, or shLPL at PID14 and maintained until PID30. For HD-LINC overexpression experiments, MSNs were transduced with lentiviruses expressing HD-LINC or LacZ control at PID14 and maintained until PID30. For ASO experiments, directly reprogrammed HD-MSNs were transfected with HD-LINC-targeting or control AP-PS ASOs using Lipofectamine RNAiMAX Transfection Reagent (Thermo Fisher Scientific, cat. no. 13778100) at PID14 and PID22. ASOs were used at a final concentration of 7.5 nM. For ferrostatin-1 treatment, directly reprogrammed HD-MSNs were treated with 5 μM ferrostatin-1 (Fer-1; MedChemExpress, HY-100579) or the corresponding DMSO vehicle control for 36 h before analysis at PID30.

### Antibodies

Primary antibodies used for immunostaining included chicken anti-TUBB3 (TUJ1; Novus, cat. no. NB100-1612, 1:1,500), rabbit anti-PLIN2/ADRP (Proteintech, cat. no. 15294-1-AP, 1:500), mouse anti-4-hydroxynonenal (4-HNE; clone 12F7; Thermo Fisher Scientific, cat. no. MA5-27570, 1:500), rabbit anti-53BP1 (Abcam, cat. no. ab21083, 1:200), mouse anti-mutant HTT (MW8; Developmental Studies Hybridoma Bank, 1:100). Alexa Fluor-conjugated goat secondary antibodies were used for fluorescent detection.

### Immunostaining and image analysis

Reprogrammed MSNs were fixed with 4% paraformaldehyde for 20 min at room temperature and permeabilized with 0.2% Triton X-100 in PBS for 10 min. Cells were blocked with 5% BSA and 1% goat serum in PBS for 1 h at room temperature and incubated with primary antibodies at 4 °C overnight, followed by fluorescent secondary antibodies for 1 h at room temperature. Cells were counterstained with DAPI and imaged using a Leica SP5X white-light laser confocal system. Imaging parameters were kept constant across experimental conditions within each experiment. Image-based quantifications were performed using FIJI/ImageJ or by manual scoring, as appropriate for the individual assay, using identical criteria across experimental conditions. Mutant HTT aggregation, 53BP1-positive signal, 4-HNE staining, PLIN2-positive lipid droplets, and neuritic degeneration were quantified as indicated in the corresponding figure legends. Neurites containing bead- or bulge-like structures were classified as dystrophic. For analysis of lipid droplets, PLIN2-positive signals were quantified separately within neuronal soma and neurites. PLIN2-positive dystrophic neurites containing detectable 4-HNE signal were scored as PLIN2-positive/4-HNE-positive neurites.

### Annexin V live-cell apoptosis assay

Live-cell apoptosis measurements were performed using the IncuCyte S3 Live-Cell Analysis System as previously described^12^. Directly reprogrammed MSNs cultured in 96-well plates were incubated with IncuCyte Annexin V Red Reagent at the indicated post-induction day. Brightfield and fluorescence images were collected at regular intervals, and Annexin V-positive objects were quantified using IncuCyte analysis software and normalized to phase area. Multiple measurements obtained from the same donor and experimental condition were averaged before statistical analysis where indicated.

### Lipid peroxidation and lipid droplet analyses

Lipid peroxidation was assessed by 4-HNE immunostaining and BODIPY 581/591 C11 staining. For live-cell BODIPY C11 measurements, directly reprogrammed MSNs were incubated with 10 μM BODIPY 581/591 C11 (Invitrogen, cat. no. D3861) for 30 min at 37 °C. Cells were washed once with PBS and imaged immediately. Oxidized and non-oxidized BODIPY C11 fluorescence was detected in the green and red channels, respectively, and lipid oxidation was quantified as the ratio of oxidized green fluorescence to non-oxidized red fluorescence. For live-cell lipid droplet imaging, directly reprogrammed HD-MSNs were labeled with BODIPY 493/503 and Hoechst 33342 and imaged for 5 min. Individual lipid droplets within neurites were tracked using FIJI, and net displacement was calculated as the distance between the initial and final position of each lipid droplet. Compartment-specific lipid droplet abundance and distribution were quantified separately within neuronal soma and neurites using identical analysis parameters across experimental conditions.

### RNA preparation and quantitative PCR with reverse transcription

Total RNA was extracted from directly reprogrammed MSNs using the RNeasy Micro Kit (Qiagen) or TRIzol Reagent (Invitrogen). Reverse transcription was performed using the SuperScript IV First-Strand Synthesis System (Invitrogen) according to the manufacturer’s protocol. Quantitative PCR was performed using SYBR Green PCR Master Mix (Applied Biosystems) and the StepOnePlus Real-Time PCR System (Applied Biosystems). Relative transcript abundance was calculated using the 2^−ΔΔCt method and normalized to the housekeeping gene HPRT1. Primer sequences were as follows: HD-LINC (LINC01139), forward 5′-GGAAGTTGATGACATTTGTAGCTG-3′ and reverse 5′-GGAAACAGGGATGCTCTGCAAAG-3′; LPL, forward 5′-CTGCTGGCATTGCAGGAAGTCT-3′ and reverse 5′-CATCAGGAGAAAGACGACTCGG-3′; and HPRT1, forward 5′-TGAGGATTTGGAAAGGGTGT-3′ and reverse 5′-AATCCAGCAGGTCAGCAAAG-3′.

### Bulk RNA sequencing and data analysis

Bulk RNA sequencing was performed on matched HD-shCon and HD-shLINC MSNs derived from four independent HD patient lines. RNA samples that passed quality control were submitted to the Genome Access Technology Center (GTAC) at Washington University in St. Louis for library preparation and sequencing. Libraries were sequenced on an Illumina NovaSeq 6000 using SMARTer 150-bp paired-end sequencing with a target depth of approximately 30 million reads per sample. Sequencing quality was assessed using FastQC. Raw FASTQ files were trimmed for adapter contamination and low-quality sequences using Cutadapt. Trimmed reads were aligned to the GRCh38 human reference genome using STAR, and raw gene counts were generated using the STAR --quantMode GeneCounts option. Raw gene counts were normalized using the trimmed mean of M-values (TMM) method, and differential gene-expression analysis between matched HD-shCon and HD-shLINC samples was performed using the exact test in edgeR. Differentially expressed genes were defined as genes with log2 fold change ≥ 0.58 or ≤ −0.58 and P < 0.05 and were used for downstream functional enrichment analyses. Weighted gene co-expression network analysis (WGCNA) was performed using normalized, transformed bulk RNA-seq expression data. Technical duplicate measurements were averaged for each biological sample, yielding four HD-shCon and four matched HD-shLINC expression profiles. Genes with no variation across samples were excluded before network construction. A signed co-expression network was generated using biweight midcorrelation, with a minimum module size of 50, deepSplit of 2, and module merge cut height of 0.30. The soft-thresholding power was selected based on scale-free topology criteria. Module eigengenes were calculated and correlated with the HD-shLINC condition using Pearson correlation. Modules with module–trait associations at P < 0.05 were considered associated with HD-LINC knockdown. Topological overlap-based networks were used to visualize representative genes within selected modules.

### Single-cell RNA sequencing and hdWGCNA analysis

Single-cell RNA sequencing was performed on directly reprogrammed MSNs derived from a pre-symptomatic HD donor and a symptomatic HD donor. Fresh cell suspensions were submitted to the Genome Technology Access Center (GTAC@MGI) at Washington University School of Medicine for library preparation using the 10x Genomics 3′ gene-expression platform. Libraries were sequenced on a NovaSeq X Plus system at a target depth of approximately 50,000 read pairs per cell. Following quality control and normalization, single-cell RNA-seq data were analyzed using Seurat in R. Dimensionality reduction was performed by principal component analysis, and cells were visualized by UMAP and clustered using the Louvain algorithm. HD-LINC expression was examined across transcriptionally defined clusters. Gene co-expression network analysis of symptomatic HD-MSNs was performed using hdWGCNA. Genes detected in at least 5% of cells in at least one symptomatic cluster were retained for network analysis. Following metacell construction, a soft-thresholding power was selected based on scale-free topology and network connectivity, and a weighted gene co-expression network was generated. Module eigengenes were calculated for each module, and module activity was additionally assessed using UCell scores. Modules were considered cluster 2-enriched when both module eigengene expression and UCell module scores were higher in cluster 2 than in each of clusters 0, 1, and 3. Intramodular connectivity was assessed using eigengene-based connectivity (kME), and genes within each module were ranked according to kME to identify representative hub genes.

### Postmortem human transcriptomic and pathway analyses

Publicly available postmortem human HD transcriptomic datasets were analyzed to determine whether transcriptional changes identified in directly reprogrammed HD-MSNs were also represented in human striatal neurons. For GSE152058, published annotations were used to identify D1 and D2 MSNs from control and HD samples. An independent dataset containing DARPP32-positive caudate neurons from control individuals and individuals with HD was analyzed in parallel. Genes identified in these datasets were compared with genes enriched in the symptomatic HD-LINC-associated neuronal state by gene-symbol overlap. Gene Ontology enrichment analysis of the shared gene sets identified across these comparisons was performed using Enrichr. Representative enriched Gene Ontology Biological Process terms were selected for visualization.

### Human brain tissue and RNAscope

All brain tissue used in the study was obtained from the New York Brain Bank. All brains were donated after consent from the next of kin or an individual with legal authority to give consent. The use of postmortem brain tissues for research was approved by the Columbia University Institutional Review Board (IRB protocol # AAAT2895). The Institutional Review Board has determined that performing clinicopathologic studies on de-identified postmortem samples is exempt from Human Subject Research according to exemption # 45 CFR 46.104(d)(2). All studies complied with ethical regulations.

Detection of LINC01139 (HD-LINC) in human postmortem formalin-fixed paraffin-embedded 7-micron thick sections was performed was done using RNAscope™ multiplex fluorescent v2 (ACDbio, Cat#323110) per the manufacturer’s protocol. Prior to proceeding, we performed a photobleaching step to eliminate autofluorescence. Slides were first baked at 60 °C for 1 h, deparaffinized in xylene with two 5-minute incubations in two fresh Xylene containers, dehydrated in 100% ethanol for twice in two separate containers, and dried at 60 °C for 5 min. The slides were then loaded into an ACD EZ-Batch holder and fully submerged in 600 mL of photobleaching buffer (1X RNase/DNase-free PBS with 0.1 mM sodium citrate, pH 7.2) in an ACD wash tray. The tray was placed inside an insulated grow tent at 4 °C in a cold room, 9 cm below a 1000–1200 W full-spectrum LED lamp with both UV and visible channels on. Slides were illuminated for 72 h, with buffer level and temperature checked every 24 h and topped up as needed. The photobleached slides were then subjected to RNAscope™ multiplex fluorescent v2 protocol as per the default instructions but without the use protease incubation step. We used predesigned ACDbio™ RNAscope probes for LINC01139 (Hs-LINC01139-C1 (REF# 506371)) and detected the signal with Opal dyes (Akoya Biosciences cat# FP1487001KT): Opal 570 at 1:1000 or Opal 690 at 1:750 dilution.

For detection of HD-LINC in cells, directly reprogrammed MSNs were cultured on coverslips until post-induction day 30 (PID30) and fixed with 4% paraformaldehyde (PFA). The coverslips were then sequentially dehydrated in 50%, 70%, and 100% ethanol for 1 min each at room temperature. Following dehydration, the coverslips were transferred to fresh 100% ethanol and stored at −20 °C until RNAscope analysis. On the day of the experiment, the cells were rehydrated and the slides were gently rinsed in PBS once at room temperature for 10 minutes, then treated with RNAscope Hydrogen Peroxide for 10 minutes at room temperature, and washed in distilled water. The cells were digested with RNAscope Protease III for 10 minutes at room temperature, followed by two washes in PBS. In situ hybridization was performed with the RNAscope Multiplex Fluorescent Reagent Kit v2 (ACD, cat. no. 323100) according to the manufacturer’s instructions. As indicated above. SNAP25 (predesigned probe, ACDbio – cat# 518851-C2) was another probe used for colabeling neurons.

For co-labeling of proteins using immunofluorescence in human brain sections, following the last step of the RNAscope protocol and prior to cover slipping, we incubated the slides with for 30 minutes at room temperature in blocking buffer (PBS + 0.3% Triton-X 100 + 1% BSA + 10% donkey serum), followed by a one-hour incubation at room temperature with the primary antibody solution (mouse anti-MAP2, Abcam, Cat# ab254143, 1:50; mouse anti-phosphorylated neurofilament heavy chain (pNF-H), Abeomics, cat34-1076, 1:20,000) in staining buffer (PBS + 0.3% Triton-X 100). This was followed by 3 washes with the RNAscope washing buffer. Next, the secondary antibody was incubated for 45 minutes at room temperature (Donkey anti-mouse, Alexa Fluor 568, Invitrogen, cat# A10037, or Donkey anti-mouse, Alexa Fluor 647, Invitrogen, cat#A-31571, 1:250). This was followed by three washes with the RNAscope washing buffer prior to adding the mounting solution, SlowFade ™ Diamond Antifade Mountant (Invitrogen, cat# S36967), and cover slipping. Tile scans images were taken on a Leica DMi98 Thunder inverted microscope 20x or 40X. For quantification of in situ hybridization scanned slides (40X magnification), the dorsal anterior caudate was annotated by a neuropathologist (OAD). We used the positive cell detection function in Qupath v0.4 to train a pixel classifier to identify all HD-LINC positive puncta, which were counted, and a density per unit area was used for statistical comparison. Brown-Forsythe ANOVA test was performed in GraphPad Prsim with post-hoc multiple comparisons and Benjamini Hochberg FDR p value correction.

### Lipidomics

Targeted lipidomic analysis was performed on matched HD-shCon and HD-shLINC MSNs derived from three independent HD patient lines. Lipid extraction and LC–MS/MS profiling were performed as previously described^30^. Briefly, lipids were extracted using a modified Bligh– Dyer method, spiked with appropriate internal standards, and analyzed using an Agilent 1260 Infinity HPLC system coupled to an Agilent 6490A triple-quadrupole mass spectrometer. Lipid species were quantified by multiple reaction monitoring in positive and negative ionization modes using corresponding internal standards. Lipid abundance was normalized to the total molar abundance of measured lipid species and expressed as molar percentage (mol%). For downstream computational analyses, mol% values were log2-transformed. For principal component analysis (PCA), each lipid species was standardized to zero mean and unit variance before PCA. Principal-component scores, explained variance, and loadings were examined to assess variation across matched HD-shCon and HD-shLINC samples and to identify lipid species contributing to treatment-associated principal components. Lipid-species-level comparisons between matched HD-shCon and HD-shLINC samples were performed using mol% values and paired two-tailed t-tests. Lipid-class-level comparisons were similarly performed between matched conditions. Heatmaps were generated from standardized lipid measurements.

### Statistics and reproducibility

Statistical analyses were performed using GraphPad Prism and R, as appropriate. Data are presented as mean ± s.e.m. unless otherwise indicated. Statistical tests and biological replicate numbers are provided in the corresponding figure legends. Paired two-tailed t-tests were used for comparisons of matched experimental conditions derived from the same donor lines where appropriate. Unpaired two-tailed t-tests were used for independent groups, and comparisons involving more than two groups were performed using one-way or two-way ANOVA followed by the indicated multiple-comparisons test. For cell-based experiments, independent patient-derived fibroblast lines were considered biological replicates unless otherwise indicated. When multiple cells, neurites, fields, or wells were measured within the same donor and experimental condition, measurements were averaged to obtain donor-level values where donor-level replication was the unit of analysis. Experiments in which individual cells or wells served as the unit of analysis are explicitly indicated in the corresponding figure legends. For lipidomic analysis, matched shCon and shLINC samples derived from the same donor were treated as paired biological samples. For postmortem tissue analyses, each independent human tissue specimen was considered one biological replicate. No statistical method was used to predetermine sample size. Sample sizes were based on the availability of independent patient-derived lines and human tissue specimens. Experimental conditions were processed in parallel with their corresponding controls.

## Data availability

Single-cell and bulk RNA-sequencing datasets generated in this study will be deposited in a public repository before publication, with accession numbers provided upon availability. Publicly available human HD transcriptomic datasets analyzed in this study are described in the Methods and cited accordingly. Lipidomic data and source data underlying the quantitative analyses will be provided with the published article. Additional data supporting the findings of this study are available from the corresponding author upon reasonable request.

## Acknowledgements

We thank Victoria Church for advice and discussions regarding live-cell and confocal imaging. We acknowledge the assistance of the Genome Technology Access Center (GTAC) at Washington University for RNA-sequencing experiments and the Biomarkers Core Laboratory at the Irving Institute for Clinical and Translational Research, Columbia University Irving Medical Center, for lipidomics profiling. Cartoons in Fig. 1A and Fig. 3A were created using BioRender.com. This study was supported by the HDF (Huntington’s Disease Foundation) Transformative Research Award (A.S.Y. and O.A.D.).

