## Supplementary Figures and Legends for "The human-specific lncRNA HD-LINC drives lipid droplet-associated neurodegeneration in Huntington’s disease"

Supplementary Figure 1

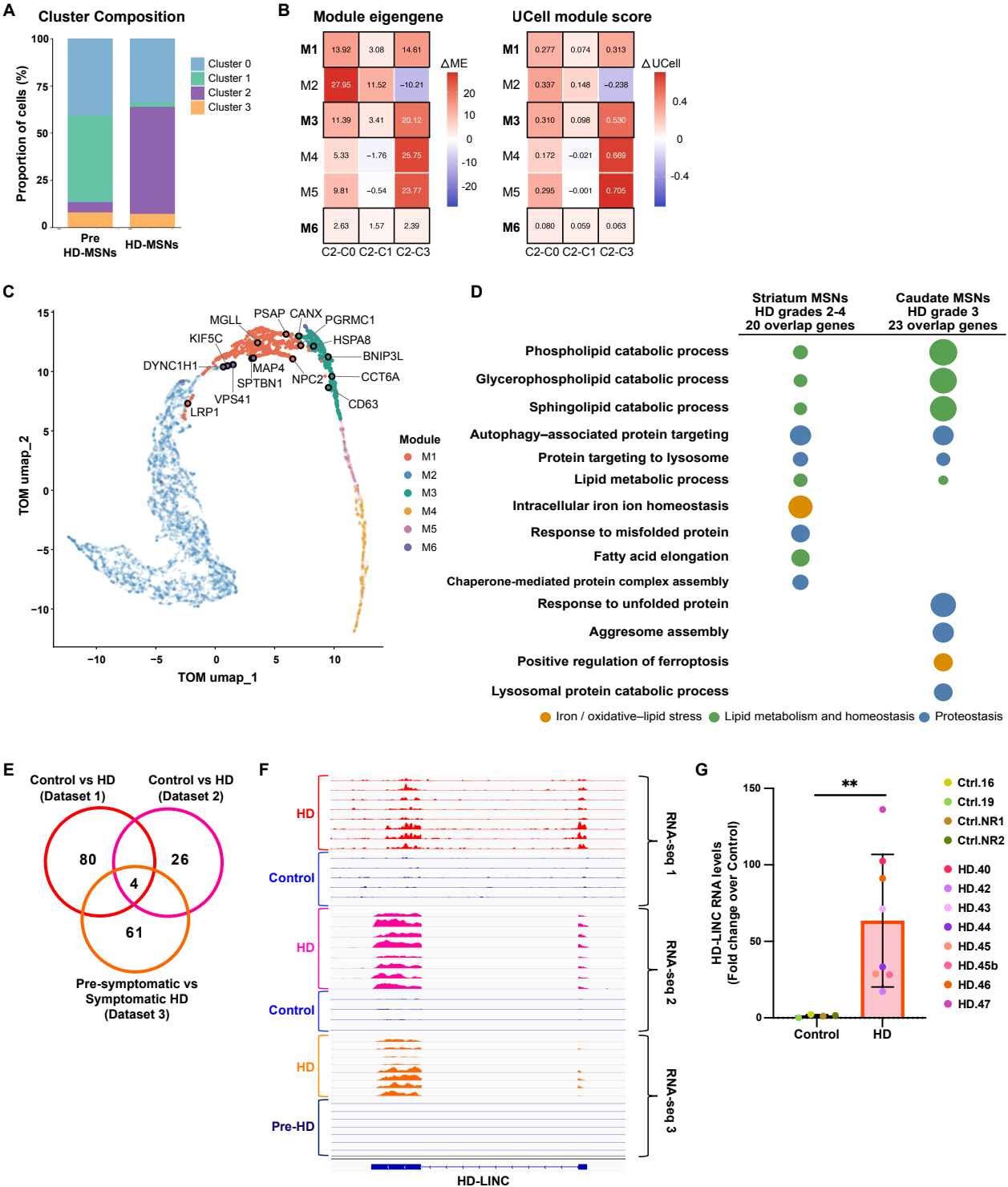

**Supplementary Figure 1. HD-LINC-associated gene networks and validation of HD-LINC expression in HD-MSNs (related to Fig. 1).**

**(A)** Relative composition of transcriptional clusters in pre-symptomatic and post-symptomatic HD-MSNs. **(B)** Heatmaps showing differences in hdWGCNA module eigengene values ($\Delta ME$; left) and UCell module scores ($\Delta UCell$; right) between the HD-LINC-enriched cluster 2 and clusters 0, 1, and 3 (C2-C0, C2-C1, and C2-C3, respectively). **(C)** TOM-based UMAP visualization of the hdWGCNA gene co-expression network in symptomatic HD-MSNs. Genes are colored according to module assignment, and selected genes are labeled. **(D)** Functional enrichment analysis of genes shared between those upregulated in the HD-LINC-enriched symptomatic HD-MSN state and those upregulated in HD relative to control neurons in two published postmortem human brain scRNA-seq datasets restricted to neuronal populations. Left, striatal D1/D2 MSNs from HD grade 2–4 samples ($n = 20$ overlapping genes). Right, caudate DARPP32-positive MSNs from HD grade 3 samples ($n = 23$ overlapping genes). Green, lipid metabolism and homeostasis; blue, protein clearance; orange, iron/oxidative-lipid stress. **(E)** Venn diagram showing significantly differentially expressed lncRNAs identified across three independent RNA-seq datasets generated from directly reprogrammed MSNs, including two control versus symptomatic HD comparisons and one pre-symptomatic versus symptomatic HD comparison. Four lncRNAs were shared across all three datasets. **(F)** Integrative Genomics Viewer (IGV) snapshots showing LINC01139 expression across the three independent RNA-seq datasets. **(G)** RT-qPCR analysis of LINC01139 expression in directly reprogrammed Ctrl-MSNs and HD-MSNs. For G, each dot represents one independent reprogrammed MSN line; $n = 4$ Ctrl-MSN lines and $n = 8$ HD-MSN lines. Statistical significance in G was determined by two-tailed unpaired t-test. Data are mean ± s.e.m. \*\*$P < 0.01$.

### Supplementary Figure 2

**A**

Human dorsal caudate (Postmortem brain)

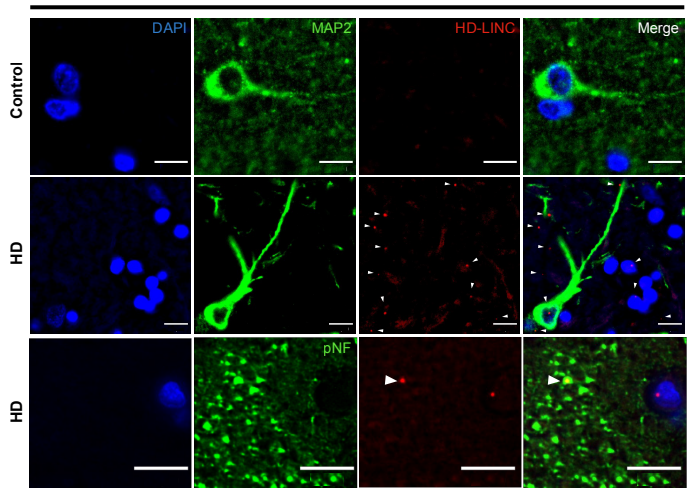

**Supplementary Figure 2. RNAscope validation of HD-LINC expression in postmortem human HD caudate (related to Fig. 1).**

**(A)** Representative RNAscope images showing HD-LINC expression in postmortem human dorsal caudate sections from control and HD samples. HD-LINC was visualized together with the neuronal markers MAP2 or phosphorylated neurofilament (pNF), as indicated. Arrowheads indicate HD-LINC puncta associated with MAP2-positive or pNF-positive neuronal structures. Nuclei were counterstained with DAPI. Scale bars, 10 μm.

Supplementary Figure 3

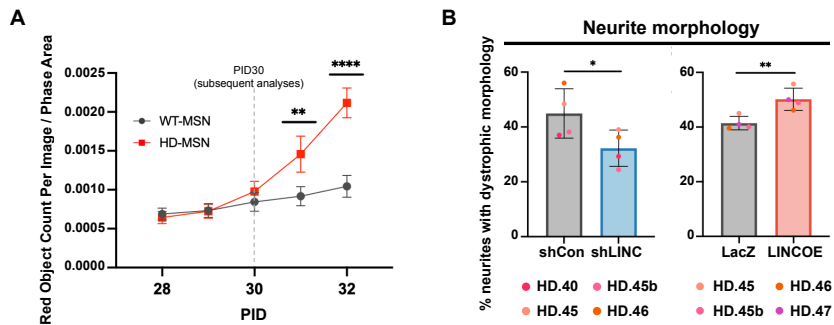

**Supplementary Figure 3. Time course of neuronal death and quantification of dystrophic neurites in HD-MSNs (related to Fig. 2).**

**(A)** Time-course quantification of Annexin V-positive red objects per image normalized to phase area in Ctrl-MSNs and HD-MSNs from post-induction day (PID) 28 to PID 32. Dashed line indicates PID 30, selected for subsequent phenotypic analyses. One Ctrl-MSN line and one HD-MSN line were analyzed, with six wells per cell line. Each plotted value in A represents the mean across the six wells. **(B)** Quantification of the percentage of neurites exhibiting dystrophic morphology in HD-MSNs following HD-LINC knockdown (left) or HD-LINC overexpression (right). Dystrophic morphology was defined as neurites containing bead- or bulge-like structures. For B, each dot represents one independent reprogrammed HD-MSN line; n = 4 HD-MSN lines per comparison. Statistical significance was determined by two-way ANOVA with Šídák's multiple-comparisons test in A and two-tailed paired t-test in B. Data are mean ± s.e.m. \*P < 0.05, \*\*P < 0.01, \*\*\*\*P < 0.0001.

Supplementary Figure 4

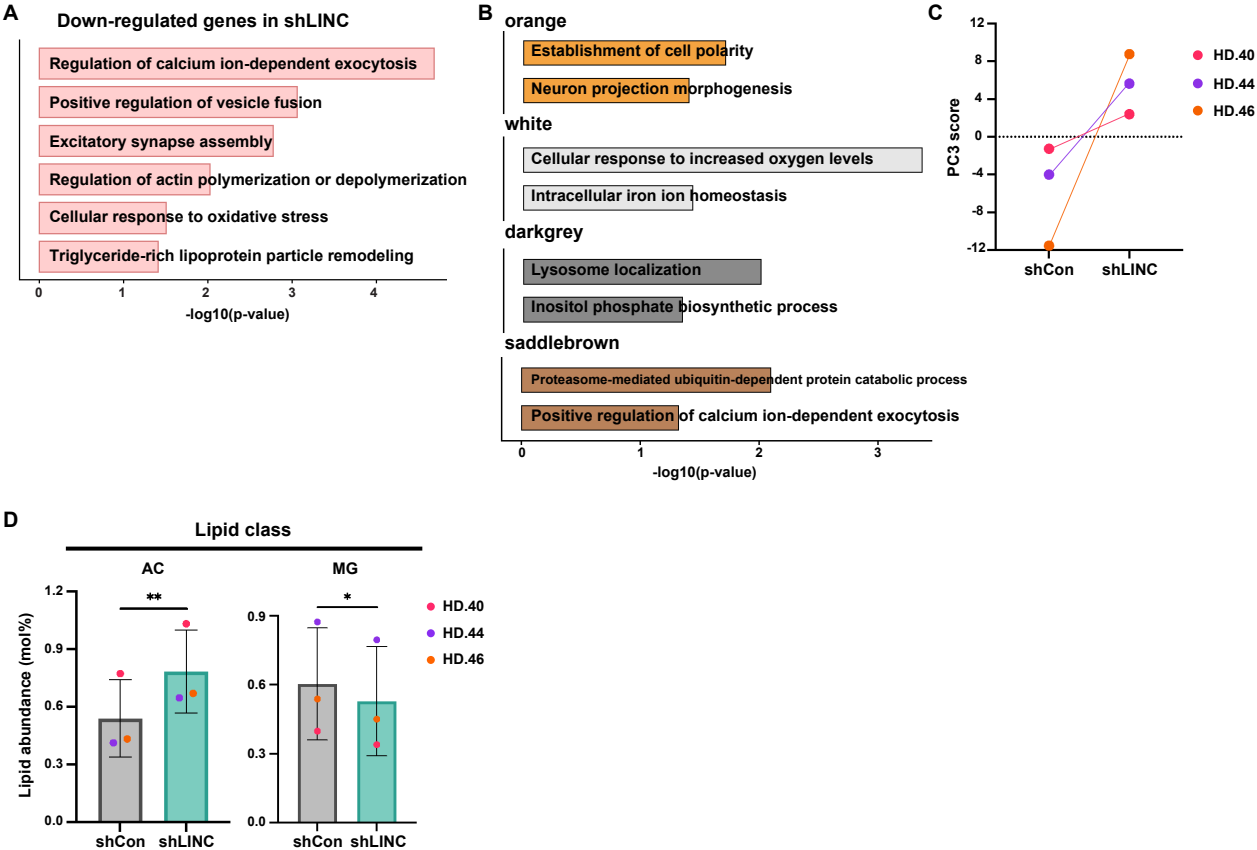

**Supplementary Figure 4. Additional transcriptomic and lipidomic changes following HD-LINC knockdown in HD-MSNs (related to Fig. 3).**

**(A)** Gene ontology (GO) terms associated with genes downregulated in HD-shLINC compared with HD-shCon MSNs. Bars represent $-\log_{10}(P\text{ value})$. **(B)** GO terms associated with the orange, white, darkgrey, and saddlebrown WGCNA modules. Bars represent $-\log_{10}(P\text{ value})$. **(C)** Paired comparison of principal component 3 (PC3) scores from targeted lipidomic profiles of HD-shCon and HD-shLINC MSNs. Samples derived from the same independent HD-MSN line are connected by lines. **(D)** Quantification of acylcarnitine (AC) and monoacylglycerol (MG) lipid-class abundance in HD-shCon and HD-shLINC MSNs. Bulk RNA-seq and WGCNA analyses in A and B were performed using $n = 4$ independent HD-MSN lines per condition. Lipidomic analyses in C and D were performed using $n = 3$ independent HD-MSN lines per condition. Each dot in C and D represents one independent HD-MSN line. Statistical significance in D was determined by two-tailed paired t-test. Data are mean ± s.e.m. \*$P < 0.05$, \*\*$P < 0.01$.

Supplementary Figure 5

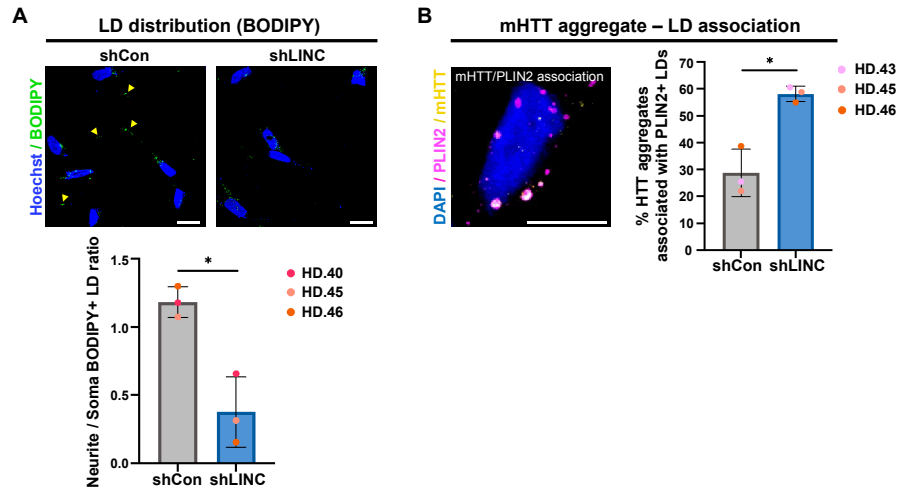

**Supplementary Figure 5. Effects of HD-LINC knockdown on lipid droplet distribution and mHTT aggregate-LD association in HD-MSNs (related to Fig. 4).**

**(A)** Representative images (top) and quantification (bottom) of the neurite-to-soma BODIPY-positive lipid droplet (LD) ratio in HD-MSNs expressing control shRNA (shCon) or shRNA targeting HD-LINC (shLINC). **(B)** Representative image (left) and quantification (right) of mutant HTT (mHTT) aggregates associated with PLIN2-positive LDs in HD-MSNs expressing shCon or shLINC. mHTT aggregates located within 1 $\mu\text{m}$ of PLIN2-positive LDs were considered LD-associated. For A and B, $n = 3$ independent HD-MSN lines per condition were analyzed, and each dot represents one independent HD-MSN line. Statistical significance was determined by two-tailed paired t-test. Data are mean ± s.e.m. \*$P < 0.05$. Scale bars, 20 $\mu\text{m}$ (Supplementary Fig. 5A) and 10 $\mu\text{m}$ (Supplementary Fig. 5B).

Supplementary Figure 6

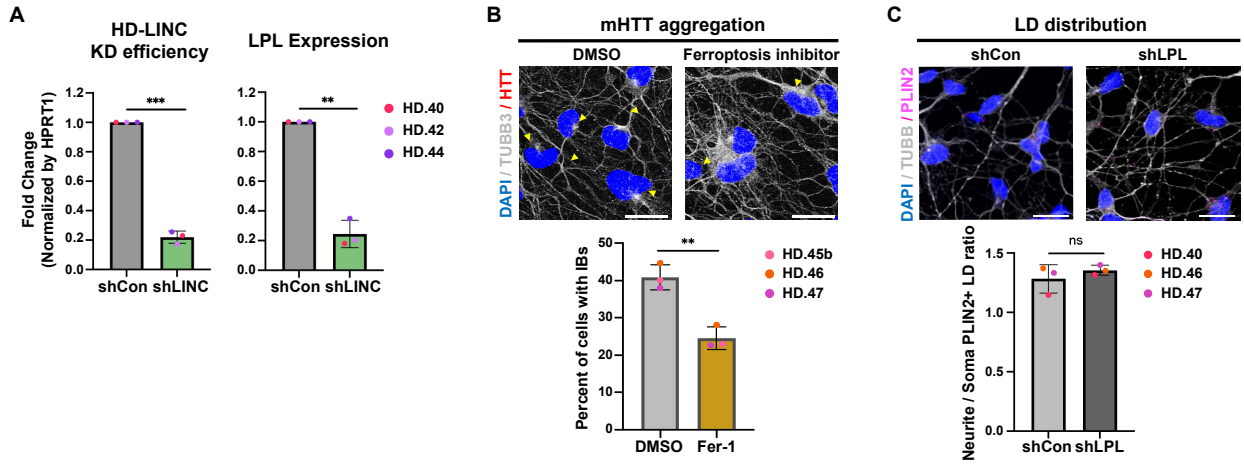

**Supplementary Figure 6. Effects of HD-LINC knockdown, ferrostatin-1 treatment, and LPL knockdown in HD-MSNs (related to Fig. 5).**

(A) RT-qPCR analysis of HD-LINC knockdown efficiency and LPL expression in HD-MSNs expressing control shRNA (shCon) or shRNA targeting HD-LINC (shLINC). Expression levels were normalized to HPRT1 and are presented relative to shCon. (B) Representative images (top) and quantification (bottom) of the percentage of HD-MSNs containing mHTT inclusion bodies (IBs) following treatment with DMSO or the ferroptosis inhibitor ferrostatin-1 (Fer-1). (C) Representative images (top) and quantification (bottom) of the neurite-to-soma PLIN2-positive lipid droplet (LD) ratio in HD-MSNs expressing shCon or shRNA targeting LPL (shLPL). For A-C, n = 3 independent HD-MSN lines per condition were analyzed, and each dot represents one independent HD-MSN line. Statistical significance was determined by two-tailed paired t-tests. Data are mean ± s.e.m. ns, not significant; \*\*P < 0.01, \*\*\*P < 0.001. Scale bars, 20 μm (B, C).

Supplementary Figure 7

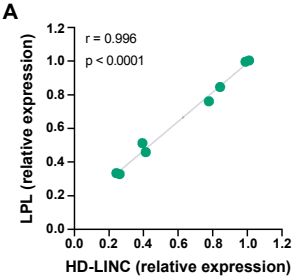

**Supplementary Figure 7. Correlation between HD-LINC and LPL expression across ASO treatment conditions in HD-MSNs (related to Fig. 6).**

**(A)** Correlation between relative HD-LINC and LPL expression following treatment with different HD-LINC ASO concentrations. Pearson's correlation coefficient ($r$) and P value are shown. One independent HD-MSN line was used, with three wells analyzed per ASO concentration. Each dot represents the paired relative expression values of HD-LINC and LPL measured in an individual well. Statistical significance was determined using a two-tailed Pearson correlation test.
